# Iterative deletion of gene trees detects extreme biases in distance-based phylogenomic coalescent analyses

**DOI:** 10.1101/2022.03.08.483551

**Authors:** John Gatesy, Daniel B. Sloan, Jessica M. Warren, Mark P. Simmons, Mark S. Springer

**Author notes:** Corresponding author. E-mail address (J. Gatesy).

## Abstract

Summary coalescent methods offer an alternative to the concatenation (supermatrix) approach for inferring phylogenetic relationships from genome-scale datasets. Given huge datasets, broad congruence between contrasting phylogenomic paradigms is often obtained, but empirical studies commonly show some well supported conflicts between concatenation and coalescence results and also between species trees estimated from alternative coalescent methods. Partitioned support indices can help arbitrate these discrepancies by pinpointing outlier loci that are unjustifiably influential at conflicting nodes. Partitioned coalescence support (PCS) recently was developed for summary coalescent methods, such as ASTRAL and MP-EST, that use the summed fits of individual gene trees to estimate the species tree. However, PCS cannot be implemented when distance-based coalescent methods (e.g., STAR, NJst, ASTRID, STEAC) are applied. Here, this deficiency is addressed by automating computation of ‘partitioned coalescent branch length’ (PCBL), a novel index that uses iterative removal of individual gene trees to assess the impact of each gene on every clade in a distance-based coalescent tree. Reanalyses of five phylogenomic datasets show that PCBL for STAR and NJst trees helps quantify the overall stability/instability of clades and clarifies disagreements with results from optimality-based coalescent analyses. PCBL scores reveal severe ‘missing taxa’, ‘apical nesting’, ‘misrooting’, and ‘basal dragdown’ biases. Contrived examples demonstrate the gross overweighting of outlier gene trees that drives these biases. Because of interrelated biases revealed by PCBL scores, caution should be exercised when using STAR and NJst, in particular when many taxa are analyzed, missing data are non-randomly distributed, and widespread gene-tree reconstruction error is suspected. Similar biases in the optimality-based coalescent method MP-EST indicate that congruence among species trees estimated via STAR, NJst, and MP-EST should not be interpreted as independent corroboration for phylogenetic relationships. Such agreements among methods instead might be due to the common defects of all three summary coalescent methods.

## 1. Introduction

Over the past two decades, phylogenetic analyses of DNA sequences increasingly have focused on genome-scale data (e.g., Rokas et al., 2003; Delsuc et al., 2005; Philippe et al., 2005; Liu et al., 2015, 2019). Despite the huge size of such datasets, many studies have documented robust conflicts between species trees generated by alternative tree-building methods. These phylogenomic conflicts include differences between topologies supported by concatenation (supermatrix) and coalescent approaches as well as between species trees estimated using alternative coalescent procedures (Zhong et al., 2013; Springer and Gatesy, 2014; Xi et al., 2014; Simmons and Gatesy, 2015; Linkem et al., 2016; Hosner et al., 2016; Simmons et al., 2016, 2019, 2022; Gatesy et al., 2017, 2019; Cloutier et al., 2019; Chan et al., 2020; Oliveros et al., 2019). The former might be expected, when incomplete lineage sorting (ILS) is a serious issue, because coalescent methods explicitly account for ILS whereas concatenation methods do not (de Queiroz and Gatesy, 2007; Kubatko et al., 2007; Edwards, 2009). By contrast, robust conflicts among species trees inferred using various ILS-aware algorithms suggest deficiencies in one or more of the coalescent approaches, even though the theoretical basis for all of these methods is similar. The multispecies coalescent with neutral sequence evolution is generally assumed (Edwards, 2009; Liu et al., 2009b, 2010; Liu and Yu, 2011; Mirarab et al., 2014; Vachaspati and Warnow, 2015).

Recent research has demonstrated that topological conflicts supported by alternative phylogenetic methods can be better understood and sometimes reconciled by partitioning support for contentious clades among the various genes that compose phylogenomic datasets (Brown and Thomson, 2017; Shen et al., 2017, 2021; Gatesy et al., 2019). Partitioned support indices initially were developed for parsimony (partitioned branch support: Baker and DeSalle, 1997; nodal dataset influence: Gatesy et al., 1999; double-decay partitioned branch support: Gatesy et al., 2003) and maximum likelihood (ML) supermatrix analyses (partitioned likelihood support: Lee and Hugall, 2003; Gatesy and Baker, 2005 [recently renamed ‘difference in gene-wise log likelihood score’ = ‘ΔGLS’ by Shen et al., 2017]; partition addition bootstrap alteration: Struck et al., 2006). An analogous Bayesian partitioned support index has shown promise for dissecting the complex patterns of among-gene support in supermatrices (Bayes factors: Brown and Thomson, 2017), and methods that summarize compatibilities/incompatibilities with independently estimated gene trees also can be used to assess the corroboration of clades supported by supermatrices or coalescent analyses (Rokas et al., 2003; Salichos and Rokas, 2013; Salichos et al., 2014; Smith et al., 2015; Kobert et al., 2016; Arcila et al., 2017).

Partitioned support indices for clades in coalescent species trees have a shorter history. Gatesy et al. (2017, 2019) presented ‘partitioned coalescence support’ (PCS), a gene-wise support index for summary coalescent methods such as MP-EST (Liu et al., 2010) and ASTRAL (Mirarab et al., 2014; Mirarab and Warnow, 2015; Zhang et al., 2017) that use optimality criteria to choose among competing species tree hypotheses. With PCS (recently renamed ‘difference in gene-wise quartet score’ = ‘ΔGQS’ by Shen et al., 2021), the support for a given clade can be divided among the constituent gene trees for a phylogenomic dataset, and the sum of all gene-tree supports for a clade, both positive and negative, equals the total support for that clade (Gatesy et al., 2019). However, this approach is not applicable for a second major class of summary coalescent methods that are commonly applied to genome-scale datasets – distance-based methods such as STAR (Liu et al., 2009b), STEAC (Liu et al., 2009b), NJst (Liu and Yu, 2011), and ASTRID (Vachaspati and Warnow, 2015). Instead of deriving phylogenomic results from the relative fits of gene trees to different species-tree topologies, these methods instead summarize average pairwise distances by counting the number of internal nodes that separate taxa in each unrooted gene tree, by specifying the rank of a shared node from the base of each rooted gene tree, or by estimating coalescence times for pairs of taxa. A phylogenetic tree is then constructed from the distance matrix using various algorithms (e.g., UPGMA, neighbor-joining, minimum evolution).

Here, we develop a simple index, ‘partitioned coalescence branch length’ (PCBL), that applies iterative removal of gene trees and repeated estimation of the species tree to quantify the influence of each gene at every supported clade of a distance-based species tree (Fig. 1). Employing the STAR and NJst methods, we automate calculation of PCBL scores for genome-scale datasets that include hundreds to thousands of loci. Phylogenomic analyses of published data from land plants (Zhong et al., 2013), spiders (Bond et al., 2014), lizards (Linkem et al., 2016), angiosperms (Xi et al., 2014), and mammals (Liu et al., 2017a) demonstrate how PCBL scores can be used to: 1) identify highly influential gene trees that drive phylogenomic conflicts, 2) determine the robustness of a species tree to removal of genes, 3) assess the impacts of missing data on summary coalescent analysis, and 4) quantify biases in distance-based phylogenomic methods. Summary coalescent methods are statistically consistent when applied to accurately reconstructed gene trees but can yield conflicting results when gene-tree reconstruction error is a factor and when a finite number of genes are sampled (e.g., Patel et al., 2013; Mirarab et al., 2016; Oliveros et al., 2019). Given that these two conditions generally apply to even the largest empirical datasets, PCBL scores are useful for arbitrating discrepancies between different coalescent methods and between concatenation and coalescent approaches. We use PCBL to characterize four severe biases that impact distance-based coalescent methods, discuss the implications of these biases within the context of biases previously documented for ASTRAL and MP-EST (Simmons and Gatesy, 2015; Gatesy et al., 2017, 2019), make recommendations on how to interpret conflicts between contrasting phylogenomic results using available support indices, and argue that several summary coalescent approaches should be used with extreme caution when many taxa are sampled, missing data are non-randomly distributed, and gene-tree reconstruction error is rampant.

**Figure 1.**
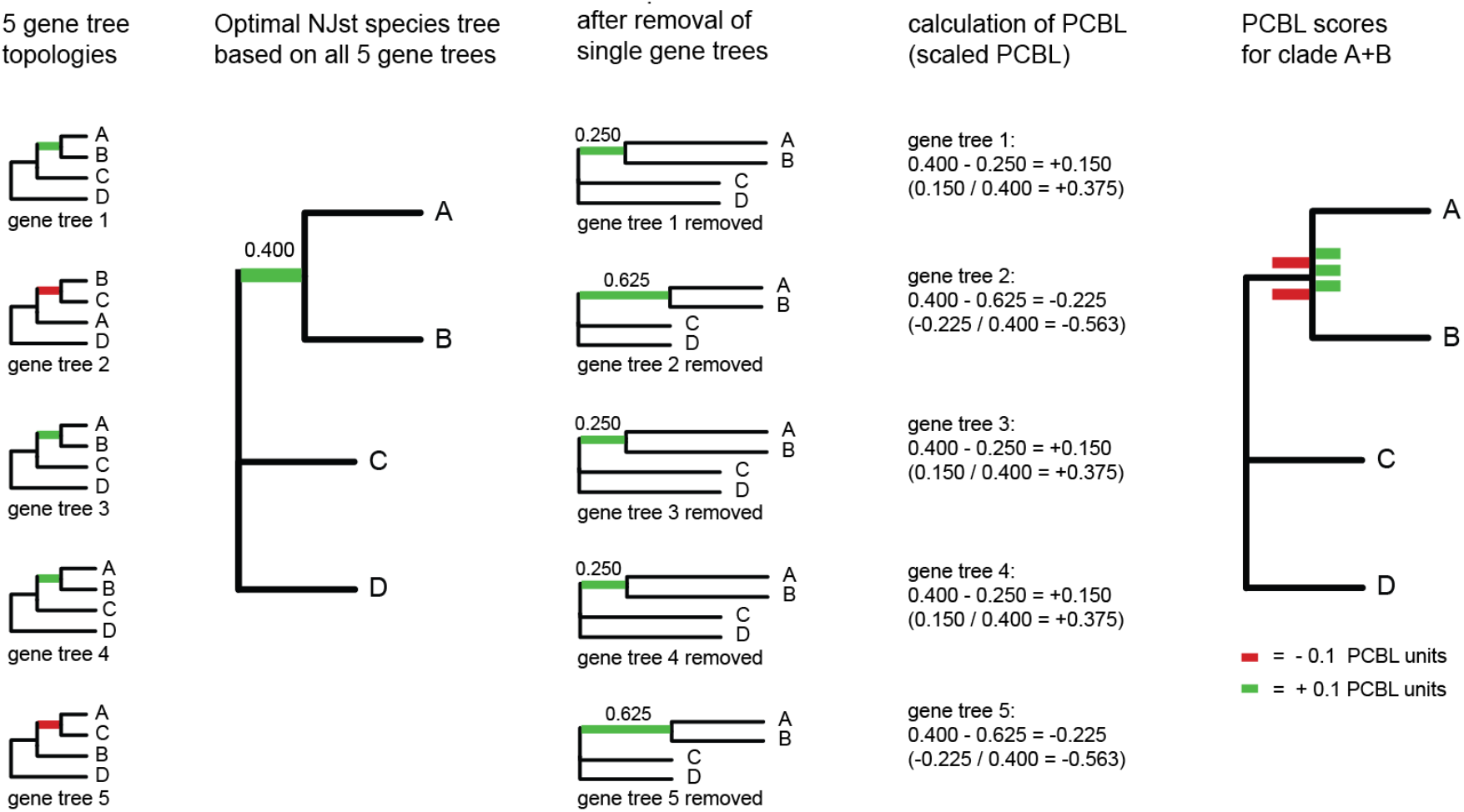
Hypothetical example that shows how partitioned coalescence branch length (PCBL) scores are calculated for five gene trees that include four taxa (A-D). PCBL for each gene tree is based on the increased or decreased length of the internal branch after that gene tree is removed from NJst analysis. The five input gene trees, the NJst species tree based on all five gene trees, NJst species trees following removal of each gene tree, calculation of PCBL (scaled PCBL in parentheses), and illustration of negative (red) and positive (green) PCBL scores for the A+B clade are shown from left to right. The same approach can be used to calculate PCBL for STAR or other distance-based coalescent methods.

## 2. Materials and Methods

### 2.1 Calculation of ‘partitioned coalescence branch length’ (PCBL) scores

For distance-based summary coalescent trees, each supported clade has a positive internal branch length. These branch lengths are derived from various pairwise distances (Liu et al., 2009b; Liu and Yu, 2011; Vachaspati and Warnow, 2013) and are distinct from the branch lengths in coalescent units that are output by MP-EST and ASTRAL. For any clade in a distance-based coalescent tree, a simple way to assess the impact of a particular gene is to delete that gene from analysis, re-estimate the species tree, and then record the impact (positive, zero, or negative) on the internal branch length that leads to the clade of interest. The difference in branch length, before and after removal of a given gene tree, is the PCBL score for that gene for that clade (Fig. 1). For any gene, the maximum possible PCBL score for a clade is equal to the original branch length that subtends the clade on the species tree. A maximum PCBL score results when removal of a single gene tree from a distance-based coalescent analysis results in the complete loss of that clade in the newly generated species tree. A smaller positive PCBL score for a clade results when removal of a particular gene tree yields a species tree that retains the clade but with a reduction in the stem branch length. A negative PCBL score is recorded when removal of a gene tree increases the branch length (Fig. 1).

By scaling PCBL to the original branch length that subtends a clade, raw PCBL scores can be expressed as proportions of the total branch length (Fig. 1). A ‘scaled PCBL’ score of +1 therefore indicates that removal of a single gene tree from analysis results in complete loss of that internal branch, while a scaled PCBL score of -1 indicates that a branch length doubles with the removal of a single gene tree from analysis. In contrast to raw PCBL scores, scaled PCBL enables a more intuitive interpretation of gene-tree influence across different internodes in the same species tree or between species trees supported by different datasets.

PCBL scores are not directly comparable to PCS scores, but both gene-wise indices measure the impact of each locus on the support for a given clade. PCS scores for a clade sum to the overall support (‘coalescence support’) for that clade, and the scaled PCS scores at a node sum to one (Gatesy et al., 2019). PCS is similar to partitioned branch support (Baker and DeSalle, 1997) and partitioned likelihood support (Lee and Hugall, 2003), which are gene-wise support indices for supermatrix analysis that are calculated by comparing the optimal species tree to the best species-tree topologies that lack a particular clade. By contrast, PCBL scores for a given internode do not sum to the overall branch length for that internode, and the scaled PCBL scores for an internode do not sum to one (Fig. 1). Removals of individual genes are used to calculate PCBL scores, and this index is therefore more analogous to ‘nodal dataset influence’, a partitioned support index for supermatrix analysis that also applies iterative removal of data partitions and re-estimation of the phylogenetic tree (Gatesy et al., 1999).

Care should be taken in interpreting PCBL scores when there is complete congruence among gene trees. If all gene trees resolve the clade of interest and there are no missing taxa in gene trees, removal of any gene tree from the analysis will affect the average pairwise distances between taxa. PCBL will be zero in such cases for every locus in the phylogenomic dataset (Fig. S1A). Furthermore, if there is complete congruence among gene trees but some taxa are missing from one or more gene trees, PCBL can be negative for a gene tree with missing taxa even though that gene tree is fully compatible with the overall species tree. In cases like this, a negative PCBL score quantifies how missing taxa can reduce the phylogenetic influence of some gene trees (Fig. S1B). PCBL simply records the impact of gene-tree removal on the internal branch lengths of a distance-based species tree (positive, negative, or neutral) and does not correct problems that certain coalescent methods have with missing taxa (Gatesy et al., 2019; this study).

### 2.2 Automation of partitioned coalescence branch length (PCBL) for genome-scale datasets

PCBL calculations were automated with custom Perl scripts (pcbl_star.pl and pcbl_njst.pl). Theses scripts take a set of gene trees as input, call the respective program (STAR or NJst as implemented in Phybase v1.5), calculate the tree based on all gene trees, iteratively remove each gene tree, re-estimate the species tree following each gene-tree removal, and then calculate PCBL for every gene at every node supported by the complete dataset of all gene trees. The scripts report when a clade is lost following the removal of any single gene tree, and all species trees (before and after each gene-tree removal) are saved to a file. For each of the five published datasets reanalyzed here (Fig. 2), PCBL scores were calculated using the STAR and NJst scripts. Code for automating PCBL analysis is freely available in the pcbl_star and pcbl_njst repositories at https://github.com/dbsloan. Given our general approach, PCBL calculations could be automated for STEAC (Liu et al., 2009b), ASTRID (Vachaspati and Warnow, 2015), or other distance-based summary coalescent methods (e.g., Mossel and Roch, 2007). We focused on NJst and STAR, because these methods have been applied previously to the datasets that were reanalyzed here.

**Figure 2.**
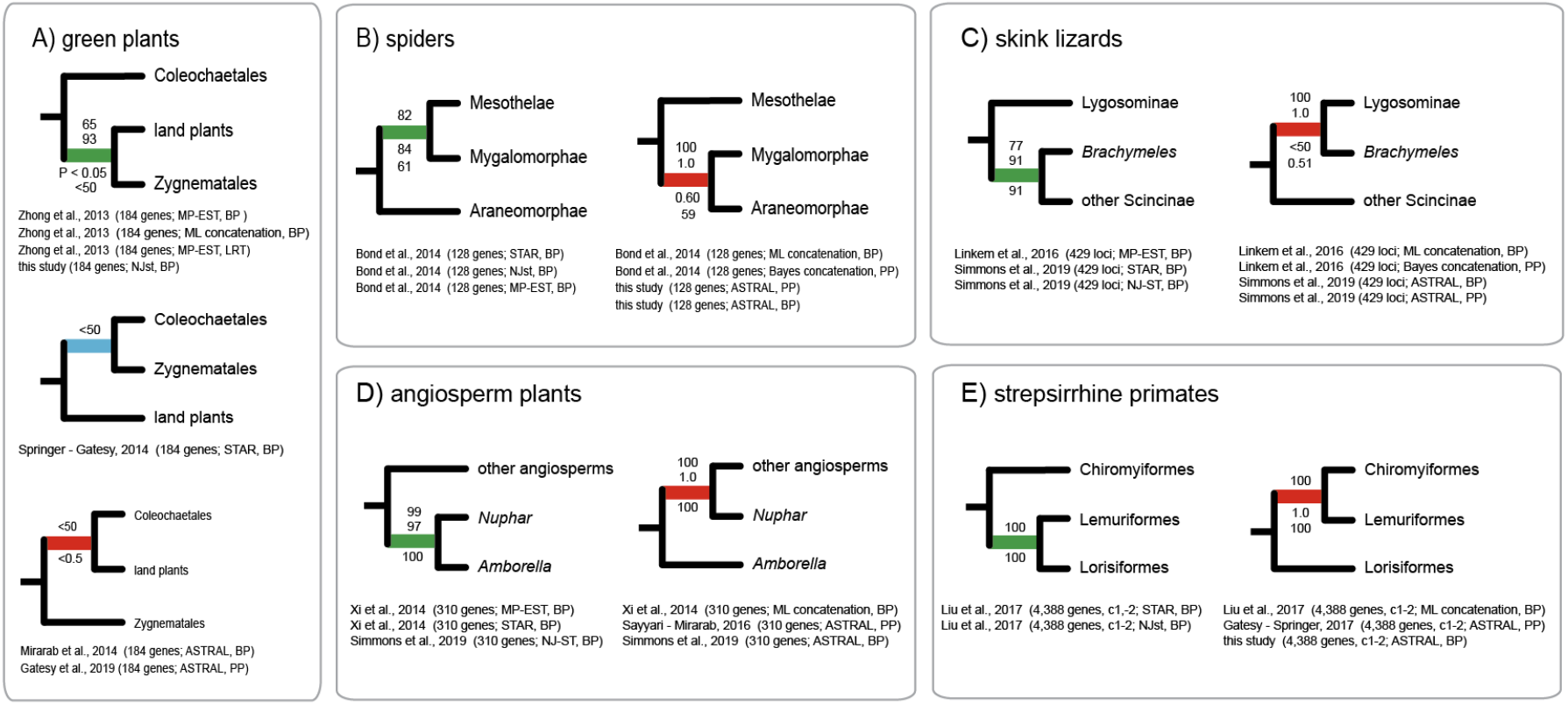
Conflicting relationships at controversial nodes for the five phylogenomic datasets reanalyzed here: A) green plants (Zhong et al., 2013), B) spiders (Bond et al., 2014), C) skink lizards (Linkem et al., 2016), D) angiosperm plants (Xi et al., 2014), and E) strepsirrhine primates (Liu et al., 2017a). For each dataset, the distance-based coalescent methods consistently conflict with concatenation and ASTRAL results. Support scores are shown above and below internal branches (likelihood ratio test = LRT, bootstrap = BP, Bayesian local posterior probability = PP). Citations for support scores are given below each tree. For E), “c1-2” indicates analysis of first and second codon positions only from protein-coding genes.

### 2.3 Published phylogenomic datasets that support conflicting resolutions at deep divergences

We reanalyzed five phylogenomic datasets (Zhong et al., 2013; Bond et al., 2014; Xi et al., 2014; Linkem et al., 2016, Liu et al., 2017a). Prior analyses of these matrices yielded conflicting relationships depending on the method of analysis, with distance-based coalescent procedures commonly contradicting ASTRAL and ML concatenation-based analyses with high support (Fig. 2). The basic features of each dataset in terms of taxa and loci that were sampled are as follows:

1. <u>Green plants</u> (Zhong et al., 2013): 184 protein-coding genes (amino acid sequences), 20 genera of land plants, nine genera of streptophyte algae, and an outgroup clade (three chlorophyte algae).
2. Spiders (Bond et al., 2014): 128 protein-coding genes (amino acid sequences), 40 ingroup taxa (spiders), and three outgroup taxa (one tick, one pseudoscorpion, one crustacean)
3. <u>Skinks</u> (Linkem et al., 2016): 429 ultraconserved element (UCE) loci (DNA sequences), 15 ingroup taxa (scincid lizards), and a single outgroup (*Xantusia*); every UCE locus was sampled for each taxon.
4. <u>Angiosperms</u> (Xi et al., 2014): 310 protein-coding genes (DNA sequences), 42 ingroup taxa (angiosperms), and four outgroup taxa (three gymnosperms, *Selaginella*).
5. <u>Mammals</u> (Liu et al., 2017a): 4,388 protein-coding genes (DNA sequences), 82 ingroup taxa (mammals), and eight outgroup taxa (two birds, one turtle, one lizard, one frog, three fishes).

### 2.4 Contrived datasets for assessing bias of high-PCBL gene trees

In addition to the core phylogenomic analyses and PCBL calculations for the five datasets listed above, the behavior of outlier gene trees (those with very high PCBL at controversial nodes) was examined further using contrived datasets. STAR and NJst species trees were inferred using implementations in Phybase v1.5 and at the STRAW website (Shaw et al., 2013; https://straw.phylolab.net/SpeciesTreeAnalysis/index.php [October, 2021]). For a given empirical dataset, an outlier gene tree at a contentious node (identified by PCBL) was combined with multiple species-tree topologies for the complete dataset. Two or more of the ‘species-tree topologies’ were combined with the single conflicting outlier gene tree to determine how strongly the outlier impacts relationships in NJst and STAR analyses. For example, we tested whether two, three, four, or many more ‘species-tree topologies’ were able to overcome the influence of a single outlier. The same analyses were executed in ASTRAL 5.6.1, a commonly used optimality-based coalescent method that is less biased than MP-EST (Simmons and Gatesy, 2015; Gatesy et al., 2017, 2019; Simmons et al., 2022).

### 2.5 Additional phylogenetic methods

Bootstrapping (Felsenstein, 1985) and Bayesian local posterior probabilities (PPs) (Sayyari and Mirarab, 2016) were used to assess support for species-tree relationships at particular nodes of interest. Bootstrap support scores for species trees generally were taken from published studies of the five datasets that were reanalyzed here (Fig. 2). When bootstrap analyses were not already published, gene-wise bootstrapping was executed using scripts from Simmons et al. (2019) for ASTRAL, STAR, and NJst (1,000 pseudoreplicates). Sayyari and Mirarab (2016) and Simmons et al. (2019) presented arguments for why gene-wise bootstrapping is preferred to site-wise bootstrapping or gene + site-wise bootstrapping. Local PPs for clades supported in maximum quartet species trees were calculated using ASTRAL 4.11.1 or ASTRAL 5.6.1. MP-EST analysis was executed for the Liu et al. (2017a) mammal dataset to provide a comparison to ASTRAL, STAR, and NJst results. One thousand independent search replicates using MP-EST v.1.5 were executed following recommendations in Simmons et al. (2016).

Sequences for selected loci with extremely high PCBL scores were examined manually to document misannotations, local misalignments, potential sequence contamination, and the extent of missing data for problematic taxa. Multiple sequence alignments were visualized using Geneious 8.1 (Kearse et al., 2012). Reciprocal BLAST searches against whole genome sequences (NCBI) were used to clarify potential homology errors (Springer and Gatesy, 2018a, 2018b). PAUP* 4.0a167 (Swofford, 2002) was used to calculate uncorrected pairwise distances between sequences as well as counts of variable and parsimony-informative sites for sequence alignments.

## 3. Results and Discussion

### 3.1 PCBL for contentious resolutions supported by distance-based coalescent methods

Our primary PCBL results are presented in five sections. For each case study, we highlight insights provided by PCBL regarding contentious relationships (in four cases, directly related to the primary phylogenetic conclusion of the published study), outlier gene trees, and severe biases in distance-based summary coalescent methods.

#### 3.1.1 Land plant origins: instability to gene-tree removal and a ‘missing taxa’ bias

For the green-plants dataset of 184 loci, MP-EST coalescent analysis supports a sister-group relationship between land plants and Zygnematales algae. Although MP-EST bootstrap support is low (65% bootstrap), likelihood ratio tests indicated high support for Zygnematales sister to land plants and rejection of alternative hypotheses at *P* < 0.05 (Zhong et al., 2013, 2014). NJst (47% bootstrap) and ML concatenation (93% bootstrap) corroborate this clade (Fig. 2A). However, STAR coalescent analysis instead supported the clade Zygnematales + Coleochaetales (44%) as the algal sister group to land plants (Springer and Gatesy, 2014), while ASTRAL produced a third topology with Coleochaetales algae as the sister to land plants (Fig. 2A; Mirarab et al., 2014; Gatesy et al., 2019).

We calculated PCBL scores for the STAR and NJst species trees (Fig. 3A, D) to assess conflicts among different coalescent methods at internodes that determine the closest living relatives of land plants (Fig. 2A), a problem that was seemingly resolved by strong likelihood ratio test support in the initial MP-EST coalescent analyses of this phylogenomic dataset (Zhong et al., 2013) but subsequently questioned upon further study (Springer and Gatesy, 2014; Gatesy et al., 2019). PCBL scores pinpointed just a few outlier gene trees that determine the alternative resolutions. For STAR (Fig. 3A-C), the removal of just one gene tree with high PCBL (#6, #146, or #178) resulted in loss of the Zygnematales + Coleochaetales clade that is sister to land plants in the original STAR tree (Fig. 3B). The NJst tree that supported Zygnematales as sister to land plants was only slightly more stable to the removal of gene trees (Fig. 3D-F). When the two gene trees with highest PCBL scores (#12 and #140) were both deleted, NJst analysis of the remaining 182 gene trees contradicted the Zygnematales plus land plants clade.

**Figure 3.**
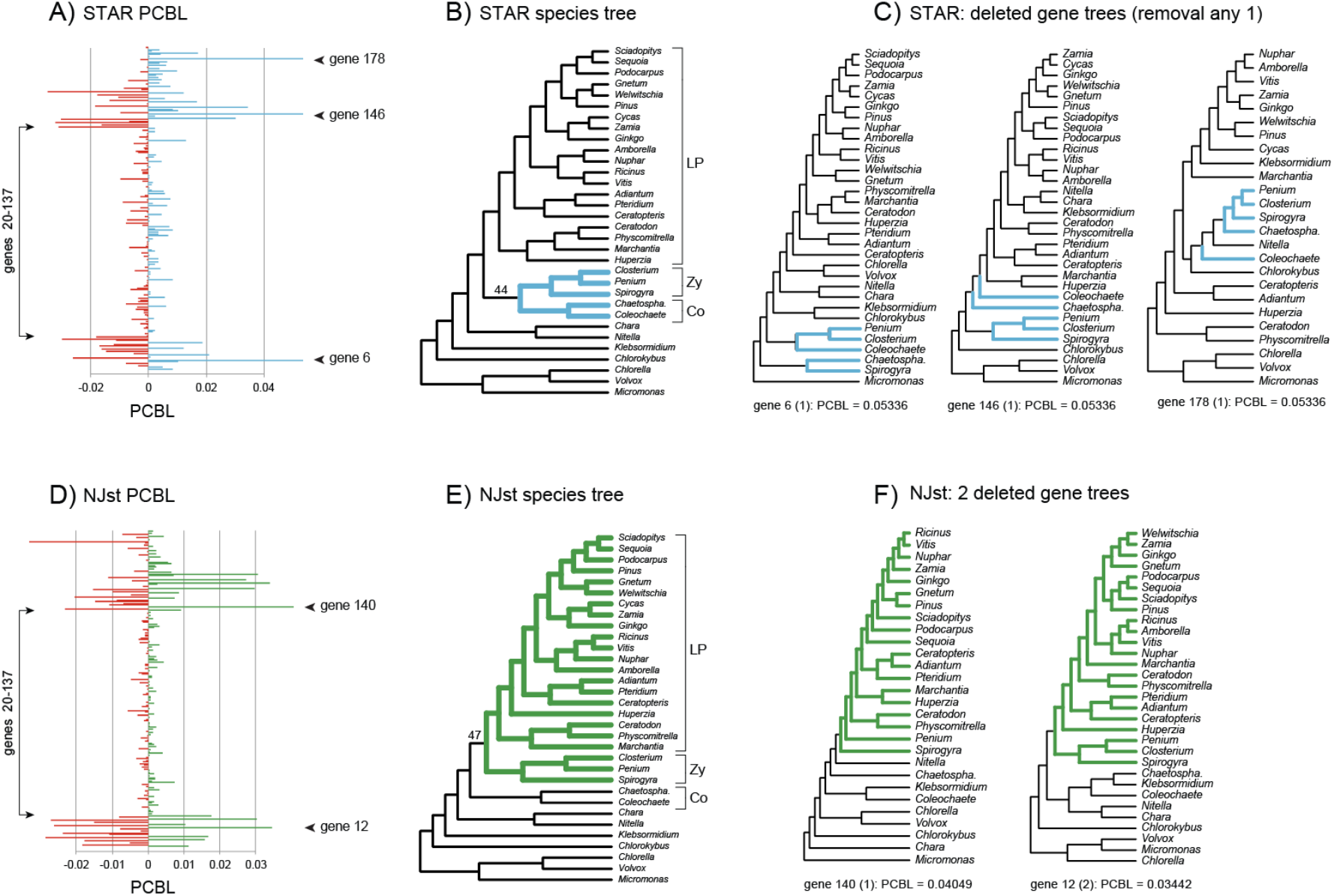
Partitioned coalescence branch length (PCBL) scores for the green-plants dataset of 184 protein-coding genes (Zhong et al., 2013) at the Zygnematales + Coleochaetales clade (STAR - blue) and the land plants + Zygnematales clade (NJst - green). For both STAR and NJst, the following are shown from left to right: PCBL scores for all gene trees (A, D), species tree based on all gene trees (B, E), and high-PCBL gene trees (C, F) that were removed to effect loss of the focal clade. For the three gene trees with highest STAR PCBL for Zygnematales + Coleochaetales, this focal clade is not resolved, but removal of these three gene trees still collapsed the Zygnematales + Coleochaetales clade. Removal of just two gene trees collapsed the land plants + Zygnematales clade supported by NJst. Small gene trees that include ≤8 taxa (genes #20-137) had uniformly low PCBL scores for both STAR and NJst. Bootstrap support for the two focal clades is shown (B, E). For each of the deleted gene trees in (C, F), PCBL score and the rank of that score (in parentheses) are shown. Abbreviations are: LP = land plants, Zy = Zygnematales, Co = Coleochaetales, *Chaetospha*. = *Chaetosphaeridium*.

For STAR, the three most influential gene trees identified by PCBL for the Zygnematales + Coleochaetales clade do not even resolve this clade. Instead, Zygnematales + Coleochaetales is poly/para-phyletic in gene trees #6, #146, and #178 (Fig. 3C). Despite not resolving the clade of interest, PCBL identified these loci as the most impactful for resolution of the Zygnematales + Coleochaetales clade. NJst PCBL analysis likewise identified gene trees that were highly influential in distance-based coalescent analysis (Fig. 3D), but in this case, the two gene trees with highest PCBL actually resolve the clade of interest, Zygnematales plus land plants (Fig. 3F).

PCBL shows how missing taxa impact the differential weighting of gene trees when summary coalescent methods are applied. Missing taxa are not randomly distributed in the green-plants dataset (Zhong et al., 2013). The same 24 taxa are absent from gene trees 20-137. These 118 small topologies with ≤ 8 taxa have consistently low PCBL scores relative to gene trees with more taxa (Fig. 3A, D; gene trees 1-19 and 138-184). Thus, as was previously demonstrated via PCS scores for MP-EST and ASTRAL (Gatesy et al., 2017, 2019), PCBL scores for STAR and NJst reveal the extreme down-weighting of small gene trees in summary coalescent analyses of genome-scale datasets. This problem (Fig. 3A, D), is clearly apparent when gene tree size (in terms of number of taxa sampled) is plotted against scaled PCBL score (online Fig. S2).

Partitioned support scores reiterate the conclusions of Springer and Gatesy (2014) and Gatesy et al. (2019) that the coalescent analyses in Zhong et al. (2013) do not robustly support Zygnematales as the sister group to land plants. Summary coalescent methods support three alternative phylogenetic resolutions with weak bootstrap support (Fig. 2A), and removals of just one or two gene trees identified by partitioned support indices disrupted relationships that define the sister group to land plants for MP-EST, ASTRAL, STAR, and NJst (Fig. 3; Gatesy et al., 2019). We therefore assert that summary coalescent analyses of this dataset have limited relevance for identifying the sister group to land plants (contra Zhong et al. 2013, 2014).

#### 3.1.2 Partial sequence for one conflicting locus determines misplacement of the segmented spider (Liphistius, Mesothelae)

Bond et al. (2014) analyzed a phylogenomic dataset comprised of 128 protein-coding loci to reconstruct the higher-level phylogeny of spiders (Araneae). A surprising conflict that emerged in their study was the unstable placement of *Liphistius* (Mesothelae; Fig. 2B), a relict genus that retains a segmented abdomen (Platnick and Sedgwick, 1984). Traditionally, *Liphistius* has been positioned as the extant sister group to remaining spiders, which include Mygalomorphae and Araneomorphae (Platnick and Gertsch, 1976). In agreement with morphological cladistics, Bayesian and ML concatenation analyses (Bond et al., 2014) grouped *Liphistius* sister to a Mygalomorphae + Araneomorphae clade (100% bootstrap, 1.0 PP). Coalescent analyses instead positioned *Liphistius* sister to Mygalomorphae (tarantulas and kin) in distance-based (82% STAR, 84% NJst) and MP-EST (61% bootstrap) trees. Our ASTRAL reanalysis of the same 128 gene trees contradicted the other coalescent methods and supported the concatenation topology with 0.60 PP and 59% bootstrap for the Mygalomorphae + Araneomorphae clade (Fig. 2B).

We used PCBL scores to examine conflicting phylogenetic placements of the mesothele *Liphistius* that were recovered with different methods (Fig. 2B). Results for NJst (Fig. 4A) and STAR (https://figshare.com/s/7cbe86e1a706514eaf35) show a similar pattern in which just a few outlier gene trees have high PCBL scores for the *Liphistius* + Mygalomorphae clade (Fig. 4B). For NJst, scaled PCBL was the maximum value (+1.0) for each of three different gene trees (#s 15, 56, 94), so removal of any one of these three gene trees overturned the controversial *Liphistius* + Mygalomorphae clade (84% bootstrap) in the original analysis of 128 genes (Bond et al., 2014). Following removal of single outliers, NJst analyses of the remaining 127 gene trees instead supported the traditional placement of *Liphistius*, in agreement with concatenation, ASTRAL, morphological evidence, and more comprehensive phylogenomic studies (Figs. 2B, 4; Platnick and Gertsch, 1976; Garrison et al., 2016; Fernández et al., 2018).

**Figure 4.**
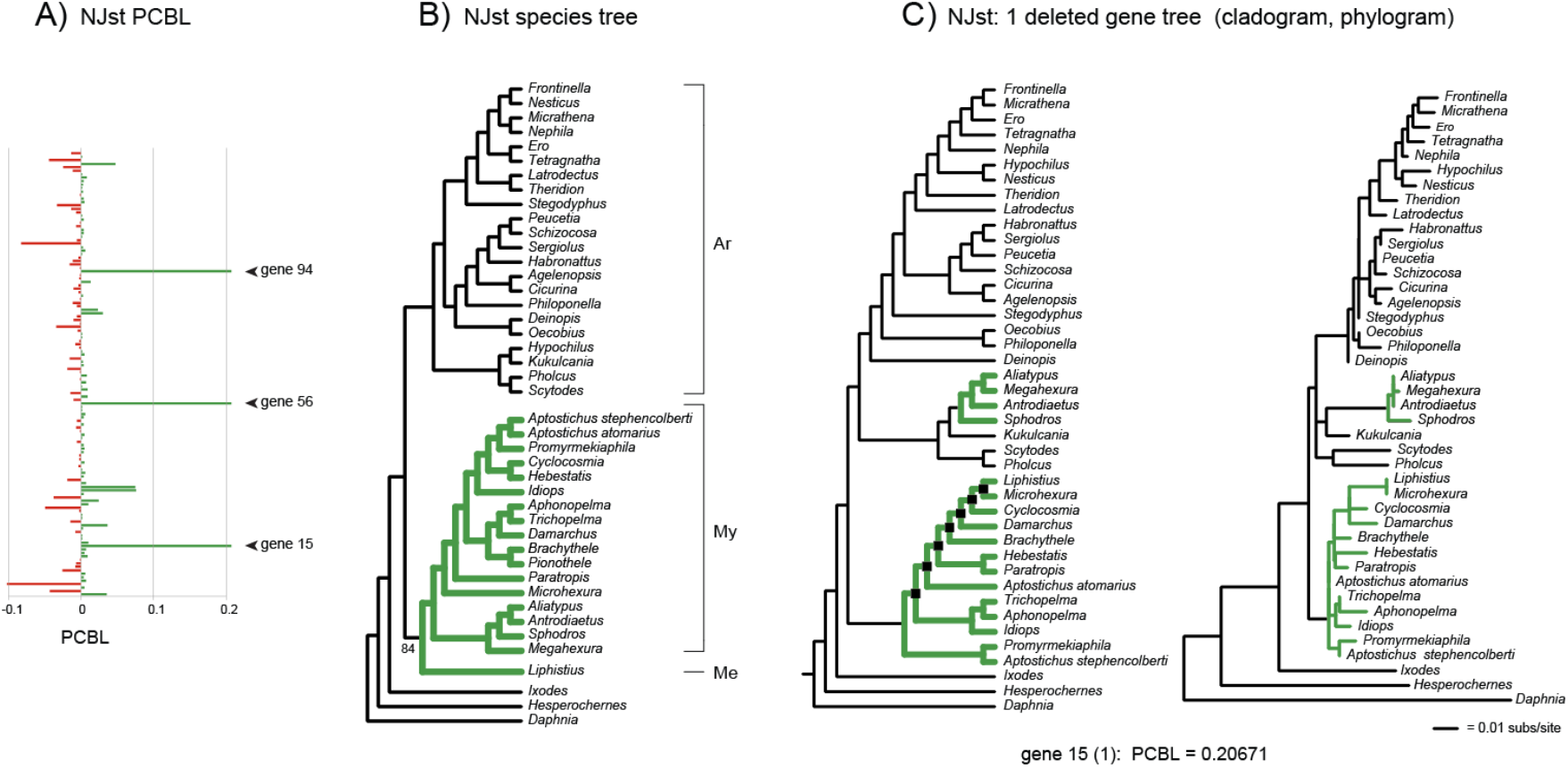
NJst partitioned coalescence branch length (PCBL) scores for the spider dataset of 128 protein-coding genes (Bond et al., 2014) at the controversial Mesothelae + Mygalomorphae clade (green). The following are shown from left to right: PCBL scores for all gene trees (A), species tree based on all gene trees (B), and one of the three gene trees with highest PCBL (C) that was removed to effect loss of the focal clade. For the three gene trees with highest PCBL for Mesothelae + Mygalomorphae (#15, #56, #94), this focal clade is not resolved, but removal of any one of these three gene trees collapsed the Mesothelae + Mygalomorphae clade. Bootstrap support for the focal clade is shown (B). For gene #15 (C), the fully resolved ML cladogram (left) and the ML phylogram (right) are shown. Black squares at internal nodes in the cladogram indicate how many nodes the mesothele *Liphistius* is nested within one subcluster of Mygalomorphae, which is polyphyletic. Note that terminal branch lengths for *Liphistius* and *Microhexura* are zero, indicating that sequences for these genera are identical. Abbreviations are: Ar = Araneomorphae, My = Mygalomorphae, Me = Mesothelae, subs/site = amino acid substitutions per site.

None of the three outlier loci with maximal NJst PCBL for the Mesothelae + Mygalomorphae clade resolve this controversial clade. Both Mygalomorphae and Araneomorphae are polyphyletic in gene tree #56, with *Liphistius* positioned within one subcluster of mygalomorphs. For gene tree #94, *Liphistius* is distant from all mygalomorphs and instead nested within a paraphyletic Araneomorphae, closest to *Pholcus, Scytodes*, and *Kukulkania*. Yet, PCBL scores correctly identified these two loci as critically influential for resolution of the *Liphistius* + Mygalomorphae clade. Presumably, the nesting of *Liphistius*, many nodes from the base of these gene trees, ‘pulled’ the divergence of this taxon up towards the apex of the species tree. The third critical locus, #15, also does not resolve the controversial *Liphistius* + Mygalomorphae clade. Mygalomorphae is polyphyletic, and *Liphistius* is nested within one of the two mygalomorph subclusters, sister to *Microhexura* (spruce-fir moss spider). The phylogram for this locus indicates no amino-acid-sequence divergence between the mesothele *Liphistius* and the mygalomorph *Microhexura* (Fig. 4C).

Given that *Liphistius* and Mygalomorphae are thought to have diverged >300 million years ago (Bond et al., 2014; Garrison et al., 2016; Fernández et al., 2018), we initially hypothesized that the sequence identity between *Liphistius* and a mygalomorph is due to contamination or a bioinformatic mixup. Examination of the multispecies alignment for locus #15 (signal recognition particle 54 [*SRP54*]) suggested a simpler explanation. *Liphistius* and several other species are represented by partial sequences, and the 103 amino acids sampled in *Liphistius* are in a conserved region of the much longer alignment (online Fig. S3). The sequence identity between very distantly related species likely represents evolutionary conservation at the amino-acid level, and it is therefore unsurprising that the short, mostly-uninformative sequence for *Liphistius* misplaced this taxon in the *SRP54* gene tree (Fig. 4C; see Lemmon et al., 2009; Hosner et al., 2016; Streicher et al., 2016 for similar examples).

#### 3.1.3 An ‘apical nesting’ bias drives conflict in phylogenomic analyses of skinks

Linkem et al. (2016) analyzed a phylogenomic dataset of 429 UCE loci from Scincidae (skink lizards). Their MP-EST coalescent analysis disagreed with their ML and Bayesian concatenation results for placement of *Brachymeles* (Fig. 2C). MP-EST (Linkem et al., 2016) and subsequent distance-based coalescent analyses (Simmons et al., 2019) placed the scincine *Brachymeles* as sister to other members of the subfamily Scincinae with 77% to 91% bootstrap support. Concatenation instead grouped *Brachymeles* sister to subfamily Lygosominae with maximum support (100% bootstrap, 1.0 PP). Linkem et al. (2016) hypothesized that multiple internodes at the base of the skink tree were in the anomaly zone (Degnan and Rosenberg, 2006) and that their MP-EST species tree was accurate. Subsequent ASTRAL coalescent analysis (Simmons et al., 2019) weakly supported the concatenation tree (Fig. 2C). Disagreements between different coalescent methods (ASTRAL vs. MP-EST, STAR, NJst) complicates Linkem et al.’s (2016) anomaly-zone interpretation and indicates that one or more of the coalescent species trees are inaccurate (Gatesy et al., 2019).

We used PCBL scores to examine further the conflicts among coalescent methods for the phylogenetic placement of *Brachymeles* (Fig. 2C). Species trees and PCBL scores for every gene tree were identical in STAR and NJst analyses (Fig. 5). For this dataset, only 12 gene trees resolve a monophyletic Scincinae, and just one (gene tree #174) places *Brachymeles* sister to remaining scincines as in the STAR/NJst/MP-EST species tree; by contrast, 33 of the 429 gene trees resolve *Brachymeles* sister to Lygosominae. PCBL scores for Scincinae were skewed strongly in the positive direction, with no PCBL scores less than -0.005 but 19 greater than +0.005 (Fig. 5A). Removal of gene trees with high PCBL scores showed that Scincinae monophyly was stable until removal of gene trees with the eight highest PCBL scores (Figure 5C). Five of the eight (#103, #205, #355, #367, #378) do not even resolve monophyly of Scincinae, including locus #378 with highest PCBL (+0.01037; Fig. 5C). Gene trees in which *Brachymeles* is highly nested within Scincinae (Fig. 5C), not more basal as in the preferred species tree (Fig. 5B), have the most impact in STAR and NJst analyses (Fig. 5A). A similar ‘apical nesting’ bias was documented in MP-EST analyses of the skink dataset when PCS was applied (figure 9 in Gatesy et al., 2019). We therefore conclude that this common bias, not problems with ML concatenation or ASTRAL, best explains the discrepancies between skink species trees supported by different phylogenomic methods.

**Figure 5.**
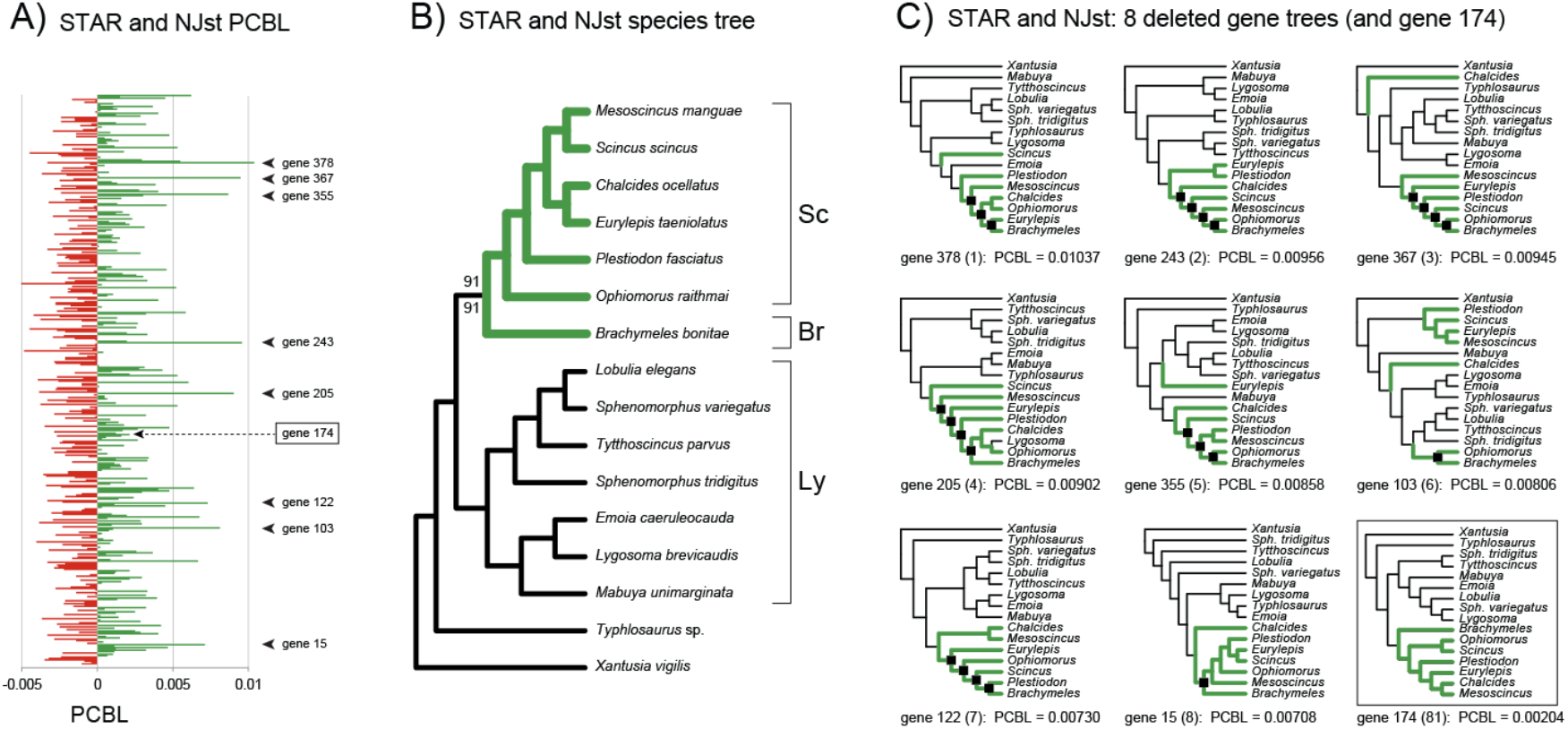
Partitioned coalescence branch length (PCBL) scores for the skink dataset of 429 UCE loci (Linkem et al., 2016) at the Scincinae clade (green). There are no missing taxa in gene trees for this dataset; species trees and PCBL scores were identical for STAR and NJst. The following are shown from left to right: PCBL scores for all gene trees (A), species tree based on all gene trees (B), and high-PCBL gene trees (C) that were removed to effect loss of the focal clade. For five of the eight gene trees with highest PCBL for Scincinae, this focal clade is not resolved. Black squares mark nodes that nest *Brachymeles* within Scincinae, instead of sister to remaining scincines as in the species tree (B). In addition to the eight gene trees with highest PCBL, the single gene tree that resolves *Brachymeles* as sister to remaining scincines is shown (C). Note the very low PCBL score for this gene tree (#174) relative to the eight outliers. STAR and NJst bootstrap support for the focal clade is shown in the species tree (B). For each of the gene trees in (C), PCBL score and the rank of that score (in parentheses) are shown. Abbreviations are: Br = *Brachymeles*, Sc = other scincines, Ly = Lygosominae, *Sph*. = *Sphenomorphus*.

#### 3.1.4 ‘Misrooting’ and ‘apical nesting’ biases distort angiosperm phylogeny

Xi et al. (2014) analyzed a phylogenomic dataset of 310 protein-coding genes from angiosperms and four outgroup taxa (Fig. 2D). ASTRAL (Mirarab and Warnow, 2015; Simmons and Gatesy, 2015) corroborated ML concatenation results (Xi et al., 2014), whereas distance-based coalescent methods (STAR, NJst) and MP-EST (Xi et al., 2014; Simmons et al., 2019) resolved an alternative topology. ASTRAL and concatenation placed *Amborella* sister to other angiosperms with high support. By contrast, distance-based coalescent methods and MP-EST grouped *Amborella* and *Nuphar* with high support and positioned this clade as sister to the remaining angiosperms. A sister group relationship between *Amborella* + *Nuphar* and the remaining angiosperms is resolved in just 28 of the 310 gene trees. By contrast, 82 gene trees resolve *Amborella* sister to all remaining angiosperms.

We used STAR and NJst PCBL scores (Fig. 6) to further clarify the striking topological differences produced by alternative phylogenomic methods (Fig. 2D). Bootstrap support for the *Amborella* + *Nuphar* clade was high for both distance-based methods (97% STAR, 100% NJst). PCBL scores for STAR and NJst showed similar patterns in which many of the same gene trees are prominent outliers. Gene trees #57, #113, #118, #157, #158, #185 are among the highest PCBL scores for both methods (Fig. 6A, D). For STAR, removal of the eight gene trees with highest PCBL scores resulted in loss of the *Amborella* + *Nuphar* clade, while the nine highest-ranking gene trees must be removed to collapse the same clade in NJst analysis (Fig. 6C).

**Figure 6.**
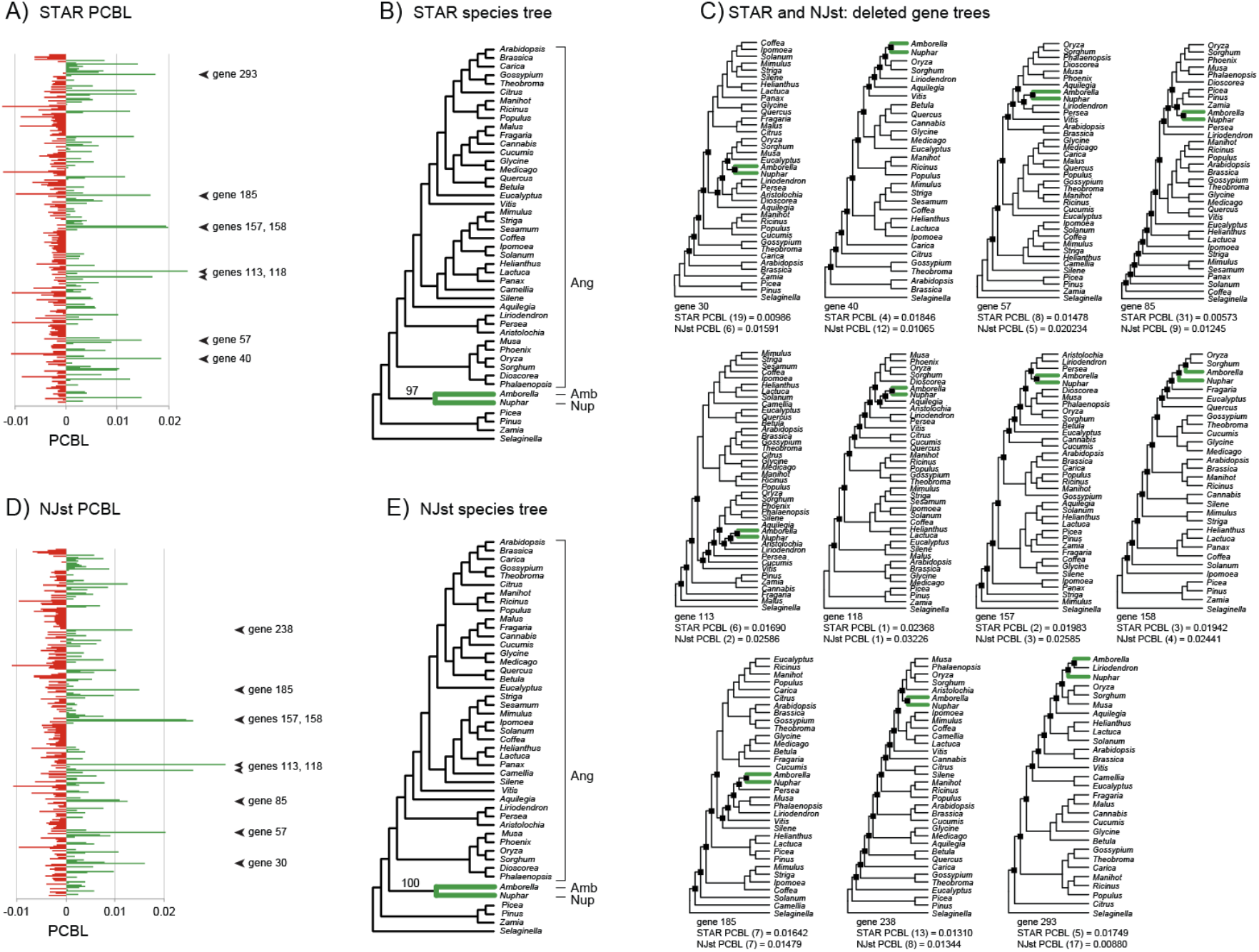
Partitioned coalescence branch length (PCBL) scores for the angiosperm-plants dataset of 310 protein-coding genes (Xi et al., 2014) at the focal *Amborella* + *Nuphar* clade (green). For both STAR and NJst, the following are shown from left to right: PCBL scores for all gene trees (A, D), species tree based on all gene trees (B, E), and high-PCBL gene trees (C) that were removed to effect loss of the focal clade. *Amborella* + *Nuphar* is not resolved in two of the 11 gene trees with high PCBL. Removal of the eight gene trees with highest STAR PCBL for *Amborella* + *Nuphar* collapsed this clade in STAR analysis; removal of the nine gene trees with highest NJst PCBL collapsed *Amborella* + *Nuphar* in NJst analysis. Bootstrap support for the focal clade is shown (B, E). For each of the deleted gene trees in (C), PCBL score and the rank of that score (in parentheses) are shown for both STAR and NJst. *Amborella* and *Nuphar* are nested multiple nodes within angiosperms (black squares at nodes) for all high-PCBL gene trees. Abbreviations are: Amb = *Amborella*, Nup = *Nuphar*, Ang = all angiosperms except *Amborella* and *Nuphar*.

The most influential gene trees in distance-based coalescent analyses according to PCBL share a common feature. *Amborella* and *Nuphar* are nested within multiple subclades of angiosperms (‘apical nesting’). Such gene trees (Fig. 6C) contradict numerous clades in the STAR and NJst species trees for this dataset (Fig. 6B and E). As noted in prior reanalyses of this dataset (Mirarab and Warnow, 2015; Simmons and Gatesy, 2015; Simmons, 2017), misrootings of the angiosperm ingroup by long-branched outgroups (Rosenfeld et al., 2012; Simmons et al, 2022) preferentially result in resolution of the *Amborella* + *Nuphar* clade, and most misroots favor a nesting of this subclade within angiosperms (see figure 2 in Simmons and Gatesy, [2015]). Two of the most influential gene trees at the *Amborella* + *Nuphar* clade that were identified by PCBL do not even resolve this clade. Gene tree #158 (3rd highest STAR PCBL; 4th highest NJst PCBL) groups *Amborella* with two grasses (*Oryza, Sorghum*), and gene tree #293 (5th highest STAR PCBL) groups *Amborella* with *Liriodendron*, the tulip tree (Fig. 6C). Again, the common theme is that *Amborella* and *Nuphar* are highly nested within angiosperms, and these outlier gene trees drive support for a clade that conflicts robustly with both ASTRAL and ML concatenation results (Fig. 2D).

#### 3.1.5 Incomplete sequences drive a ‘basal dragdown’ of the aye-aye (Daubentonia) in strepsirrhine primate phylogeny

Liu et al. (2017a) analyzed a phylogenomic dataset of 4,388 genes from mammals. We reanalyzed their gene trees based on first and second codon positions from protein-coding loci (Liu et al., 2017a; their figure 1). Phylogenomic results for these data (Figs. 2E, 7C) revealed several strongly supported conflicts between ML concatenation and distance-based coalescent methods (STAR, NJst). Subsequent reanalysis indicated that the species tree supported by ASTRAL conflicted robustly with both distance-based coalescent methods and instead agreed with ML concatenation at several nodes (Gatesy and Springer, 2017; this study). One of the most surprising conflicts in the distance-based coalescent trees involved placement of the aye-aye, *Daubentonia* (Chiromyiformes), a highly specialized strepsirrhine primate that is well established as the sister group to lemurs (Lemuriformes). This sister-group relationship is supported by mitochondrial genomes (Finstermeier et al., 2013), supermatrices (Fabre et al., 2009; Perelman et al., 2011; Springer et al., 2012), nuclear genomes (Marciniak et al. 2021), and low-homoplasy retroelement insertions (McLain et al., 2012). By contrast, STAR and NJst (Fig. 2E) positioned *Daubentonia* as sister to a well-supported clade (100% bootstrap) composed of *Otolemur* (Lorisiformes) and *Microcebus* (Lemuriformes).

We calculated STAR and NJst PCBL scores to critically examine the suspicious support for the *Otolemur* + *Microcebus* clade within Strepsirrhini (Fig. 7). Species trees for STAR and NJst were topologically identical (Fig. 7C) and show similar PCBL profiles with strong outliers in the positive direction for the grouping of *Otolemur* + *Microcebus* (Fig. 7A-B). In the four gene trees with highest PCBL (#86, #878, #1031, #2717), *Daubentonia* is widely separated from its likely sister group, *Microcebus*, by 26-34 internal nodes (Fig. 8), and most clades in these gene trees conflict with the overall distance-based species tree (Fig. 7C). Gene tree #86 does not even resolve the *Otolemur* + *Microcebus* clade but, according to PCBL scores, is the 2nd (STAR) or 3rd (NJst) most influential gene tree at this node.

**Figure 7.**
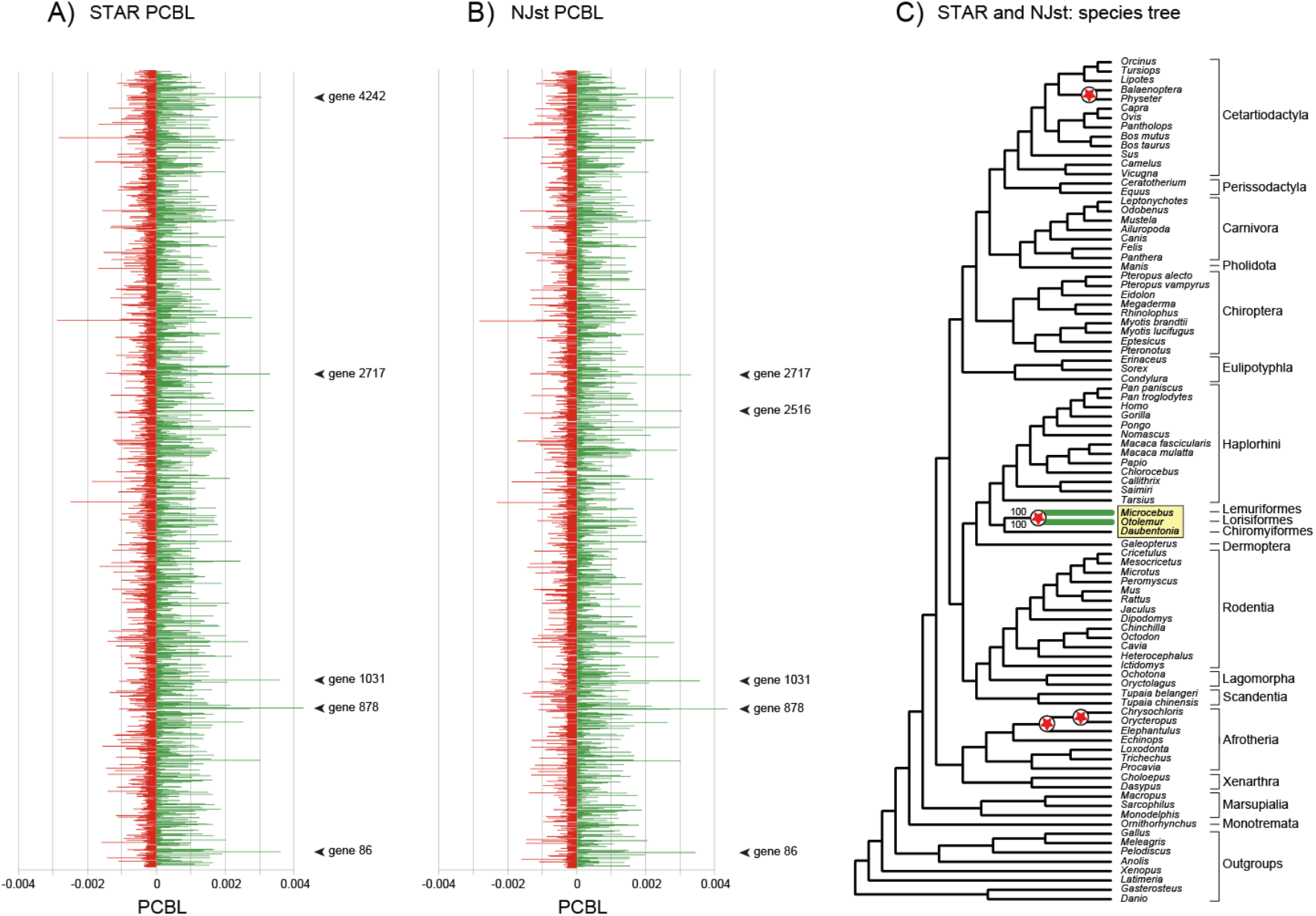
Partitioned coalescence branch length (PCBL) scores for the mammal dataset of 4,388 protein-coding genes (Liu et al., 2017a) at the Lemuriformes + Lorisiformes clade (green). The following are shown: STAR PCBL (A), NJst PCBL (B), and the species tree based on all gene trees for both STAR and NJst (C). Arrowheads mark gene trees with the five highest PCBL scores in STAR (A) and NJst (B) analyses. In the species tree (C), strepsirrhine primate genera are enclosed in a yellow rectangle, and bootstrap support scores for the dubious focal clade is indicated for STAR and NJst. Red stars at internal nodes mark the four unconventional clades robustly supported by the distance-based coalescent methods (100% bootstrap) that strongly conflict with ASTRAL analysis of the same data. ASTRAL instead supported clades favored by previous large-scale phylogenetic analyses with robust support (bootstrap = 98-100%; PP = 0.99-1.0). Higher-level mammalian taxa and outgroups are delimited to the right of genus names.

**Figure 8.**
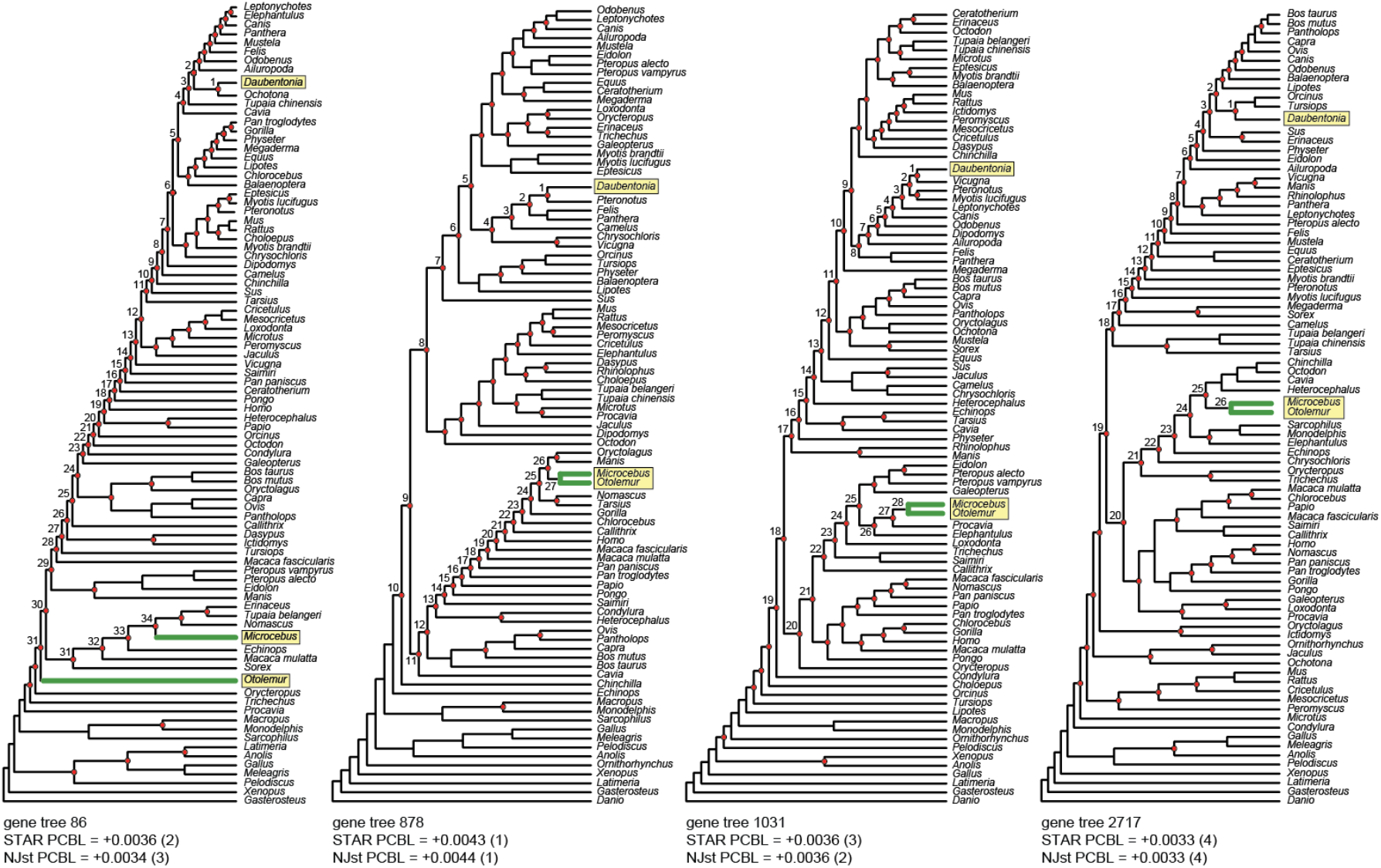
High-PCBL gene trees at the dubious Lemuriformes + Lorisiformes clade (green) for the mammal dataset of 4,388 protein-coding genes (Liu et al., 2017a). PCBL score and the rank of that score (in parentheses) are shown for both STAR and NJst. Red circles at nodes mark the numerous clades that conflict with the distance-based species tree (Fig. 7C). Strepsirrhine primate genera are enclosed in yellow rectangles. Note that the aye-aye (*Daubentonia* - Chiromyiformes) is widely separated from *Microcebus* (Lemuriformes) by many internal nodes in each gene tree. These internal nodes are numbered in the four high-PCBL gene trees. The second (STAR) or third (NJst) highest PCBL score was recorded for gene tree #86 that does not even resolve the Lemuriformes + Lorisiformes clade.

*Daubentonia* is grossly misplaced in all of these high PCBL gene trees, and in each case, the aye-aye does not group with other primates (Fig. 8). *Daubentonia* is instead sister to a pika (*Ochotona* for gene tree #86), to a bat (*Pteronotus*; #878), to a camel (*Vicugna*; #1031), or to a clade of two dolphins (*Tursiops* + *Orcinus*; #2717). Indeed, examination of gene trees with the 20 highest PCBL scores for the controversial *Otolemur* + *Microcebus* clade revealed similar patterns. In every case, *Daubentonia* is widely separated by many internodes from the other two strepsirrhine primates, and the *Otolemur* + *Microcebus* clade is not even resolved in many of the gene trees that exert extremely high influence at this node. Even when gene trees with the 50 highest PCBL scores were deleted, STAR still supported the aberrant *Otolemur* + *Microcebus* clade, which further indicates the strength of the biased support for this grouping.

Prior scrutiny of the mammal dataset (Gatesy and Springer, 2017; Liu et al., 2017b; Wu et al., 2018; Du et al., 2019) revealed widespread homology errors in sequence alignments due to misannotation of introns and other non-coding regions in combination with misalignment of these regions to exons in the protein-coding dataset of Liu et al. (2017a). We initially hypothesized that these ‘non-coding contamination’ problems drove the high support for *Otolemur* + *Microcebus* and other unconventional mammalian clades (Fig. 7C; Gatesy and Springer, 2017). Such homology errors are present in many of the protein-coding alignments that yielded gene trees with high PCBL, but these mistakes generally impacted other clades supported by the species tree, not the *Otolemur* + *Microcebus* grouping (e.g., gene #878; online Fig. S4).

Scrutiny of DNA alignments for genes with the highest PCBL for *Otolemur* + *Microcebus* generally showed that incomplete sequences for *Daubentonia* can explain the wide separation of this taxon from other strepsirrhine primates (Fig. 8). For example, the gene tree with highest PCBL for both STAR and NJst (#878; *JOSD1*) was derived from an alignment that is 597 bp in length, but the *Daubentonia* sequence includes just 46 bp. Because this sequence is so short, it is indistinguishable from four distant mammalian relatives in the multi-species alignment for this locus (Online Fig. S4). Other high-PCBL genes showed similar patterns with very limited sequence information for *Daubentonia*. For gene #86 (*AP2M1*), the *Daubentonia* sequence is just 538 bp, and there are only 26 parsimony-informative sites for the 80 mammalian taxa in the overall alignment of 1,106 bp. *Daubentonia* shows no sequence divergence relative to 39 other taxa in the *AP2M1* alignment, which explains the highly conflicting gene tree (Fig. 8) that is almost wholly incongruent with the distance-based species tree (Fig. 7C) due to arbitrary resolution of many unsupported clades in the fully-resolved ML gene tree (Sayyari et al., 2017; Simmons and Gatesy, 2021). For the 90 taxa in the mammal dataset, *Daubentonia* is characterized by the most missing data; 2,607,345 bp were sampled from 3,721 loci (701 bp per gene). By contrast, for the remaining taxa in this dataset, an average of 6,781,709 bp were included from an average of 4,210 loci (1,606 bp per gene). Most high PCBL genes at the controversial *Otolemur* + *Microcebus* node are characterized by extensive missing data for *Daubentonia* and/or few informative sites (see https://figshare.com/s/7cbe86e1a706514eaf35 for PCBL scores and Liu et al., 2017a for DNA sequence alignments).

More than 2.6 million bp could be interpreted as an impressive sample of systematic data. But extensive *missing* sequence for *Daubentonia* relative to other taxa, in combination with insufficient informative variation in protein-coding alignments that excluded third codon positions, resulted in the gross misplacement of this genus in ML gene trees (e.g., Fig. 8) with downstream distortion of distance-based species trees (Fig. 7C). Inaccurate, highly-influential, outlier gene trees overwhelmed support for the traditional clade *Daubentonia* + *Microcebus* and dragged *Daubentonia* toward the base of the tree, away from its well-corroborated phylogenetic position that is supported by both ASTRAL and ML concatenation (100% bootstrap, 1.0 PP).

### 3.2 Phylogenetic impacts of outlier gene trees identified by high PCBL scores

In a recent review of ‘modern phylogenomics’, Liu et al. (2019: p. 227) asserted that, “One attractive prospect of algorithms for species tree construction that use summary statistics, such as STAR and STEAC, is that these methods are powerful and fast, yet they appear less susceptible to error due to deviations of single genes from neutral expectations. These methods do not utilize all the information in the data and hence can be less efficient than Bayesian or likelihood methods [Xu and Yang, 2016], yet they perform well with moderate amounts of gene tree outliers…”

This assertion has not, however, been assessed critically in empirical datasets, because indices for identifying the most influential genes at highly controversial nodes were lacking for distance-based coalescent methods. As shown below, simple contrived examples based on our PCBL analyses of empirical datasets show how extreme the biased overweighting of single outlier gene trees can be in distance-based coalescent methods.

#### 3.2.1 ‘Missing taxa’ bias in distance-based coalescent methods

We first assessed the impacts of missing taxa that were evident in the green-plants dataset (Zhong et al., 2013). Figure 9 shows a hypothetical case in which a very small gene tree (#107) with just six taxa was combined with a large gene tree that includes all 32 taxa in the dataset and matches the STAR species-tree topology (Fig. 3B). The STAR tree resolves the Zygnematales + Coleochaetales clade as sister to land plants. The small gene tree conflicts at just one node with the larger tree and instead resolves Coleochaetales (represented by *Coleochaete*) as the sister group to land plants (represented by *Physocomitrella*), which is the ASTRAL resolution (Fig. 2A). Even when 100 of the small gene trees were combined with just one large tree, STAR and NJst analyses supported a species tree that exactly matches the single large input tree which supports Zygnematales + Coleochaetales (Fig. 9). This inferred species tree contradicts 100 of the input gene trees and implies 100 deep coalescences, one for each small gene tree (Fig. 4), whereas a species tree that supports Coleochaetales sister to land plants would imply just a single deep coalescence. Even when 1,000,000 of the small gene trees were combined with the single large tree, the same distance-based species tree was inferred. This example clearly demonstrates the distortions due to missing taxa for the green-plants dataset wherein small gene trees have generally low impact relative to larger gene trees (Fig. 3A, D; online Fig. S2).

**Figure 9.**
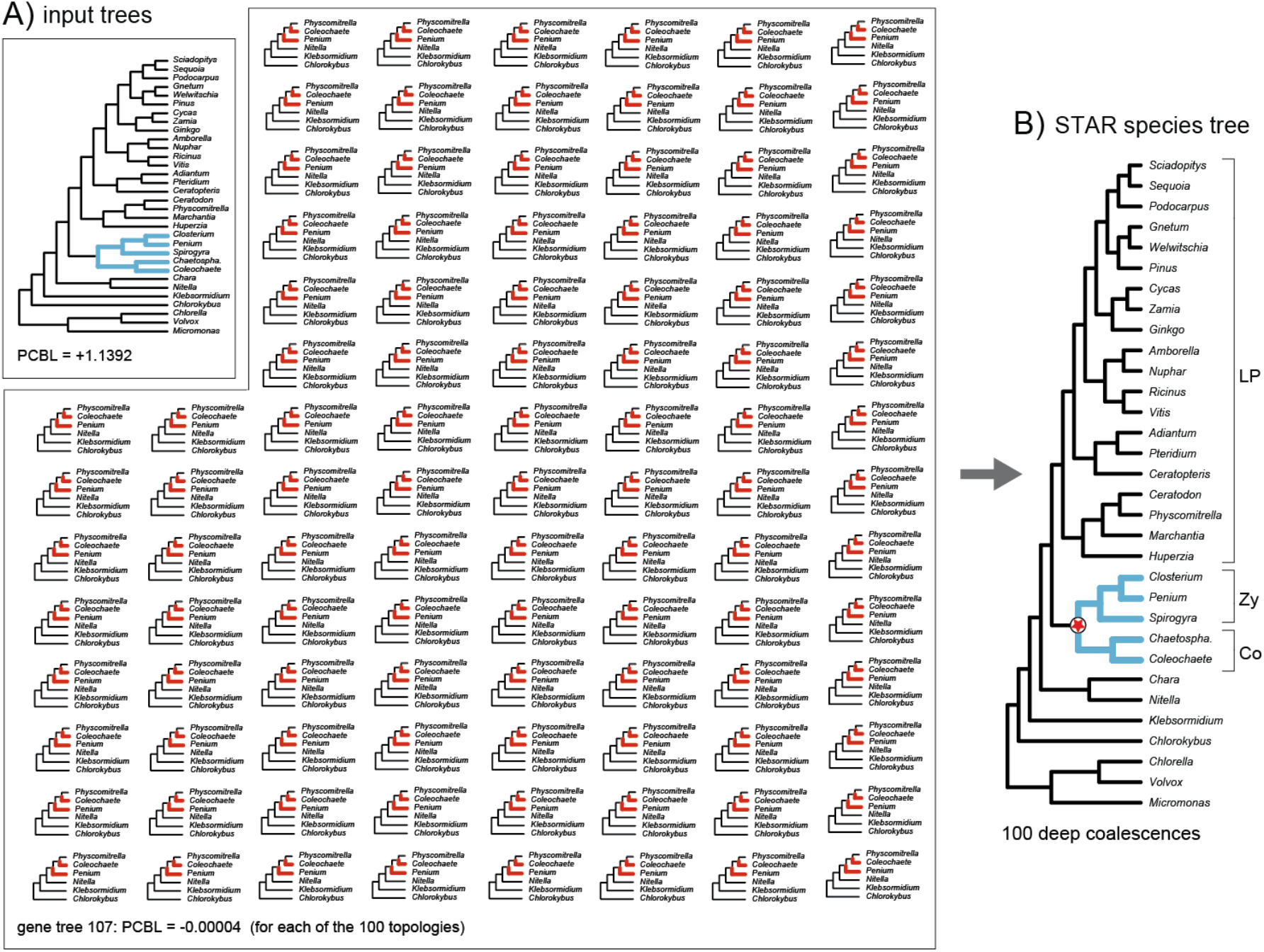
A hypothetical example that demonstrates the extreme impact of missing taxa in gene trees (A) for inference of a distance-based species tree (B). STAR analysis of 100 identical small gene trees (six taxa each) and a single conflicting larger tree (32 taxa) yielded a species tree that matched the single large input tree and contradicted the 100 small gene trees. The output species tree implies minimally 100 deep coalescences (one for each small gene tree), which is 99 more deep coalescences than is necessary. The red star (B) marks the single clade in the species tree that conflicts with each of the small gene trees (A). The large input topology with 32 taxa matches the STAR species tree for the full green-plants dataset (Zhong et al., 2013), and the 100 small input trees match gene tree #107.

A similar bias was characterized for both MP-EST and ASTRAL in which small gene trees had very little weight in analysis relative to larger gene trees (Gatesy et al., 2019, their figures 6 and 12). For the commonly used ASTRAL method, the contrived scenario illustrated in Figure 9 revealed that 280 of the small gene trees overturned the single large gene tree to yield a species tree that supports Coleochaetales sister to land plants (just one deep coalescence). However, 279 identical small gene trees combined with a single large gene tree yielded a species tree that supports Zygnematales + Coleochaetales as in the single large gene tree (279 deep coalescences). This is a reminder that the ‘missing taxa’ bias is a potentially severe problem for both distance-(NJst, STAR) and optimality-based (ASTRAL, MP-EST) coalescent methods (Fig. S2).

#### 3.2.2 ‘Apical nesting’ and ‘misrooting’ biases in distance-based coalescent methods

For the angiosperm-plants dataset (Xi et al., 2014), outlier gene trees with high PCBL scores at the controversial *Amborella* + *Nuphar* node consistently expressed ‘misrooting’ and ‘apical nesting’ biases (Fig. 6A, D). Figure 10 shows a hypothetical case in which seven perfectly congruent topologies that match the ASTRAL species tree for this dataset (*Amborella* sister to remaining angiosperms) were combined with a single conflicting gene tree that resolves the *Amborella* + *Nuphar* clade (Fig. 10). This outlier gene tree (#118) had the highest PCBL score for the contentious *Amborella* + *Nuphar* clade in both STAR and NJst analyses (Fig. 6). When the single gene tree that resolves the *Amborella* + *Nuphar* clade nested 12 nodes within angiosperms was combined with seven identical gene trees that contradict this clade, the *Amborella* + *Nuphar* clade was supported by STAR (Fig. 9). The inferred STAR species tree implies seven deep coalescences in the seven perfectly congruent input trees (Fig. 10). Results are similar for NJst. When six of the ASTRAL species tree topologies were combined with the single outlier (gene tree #118), the *Amborella* + *Nuphar* clade was again supported, even though six of the seven input trees directly contradict this clade. PCBL scores (Fig. 6) correctly predicted the biased influence of this outlier gene tree. The commonly used coalescent method ASTRAL does not suffer from this bias in which a single outlier overturns multiple perfectly congruent trees.

**Figure 10.**
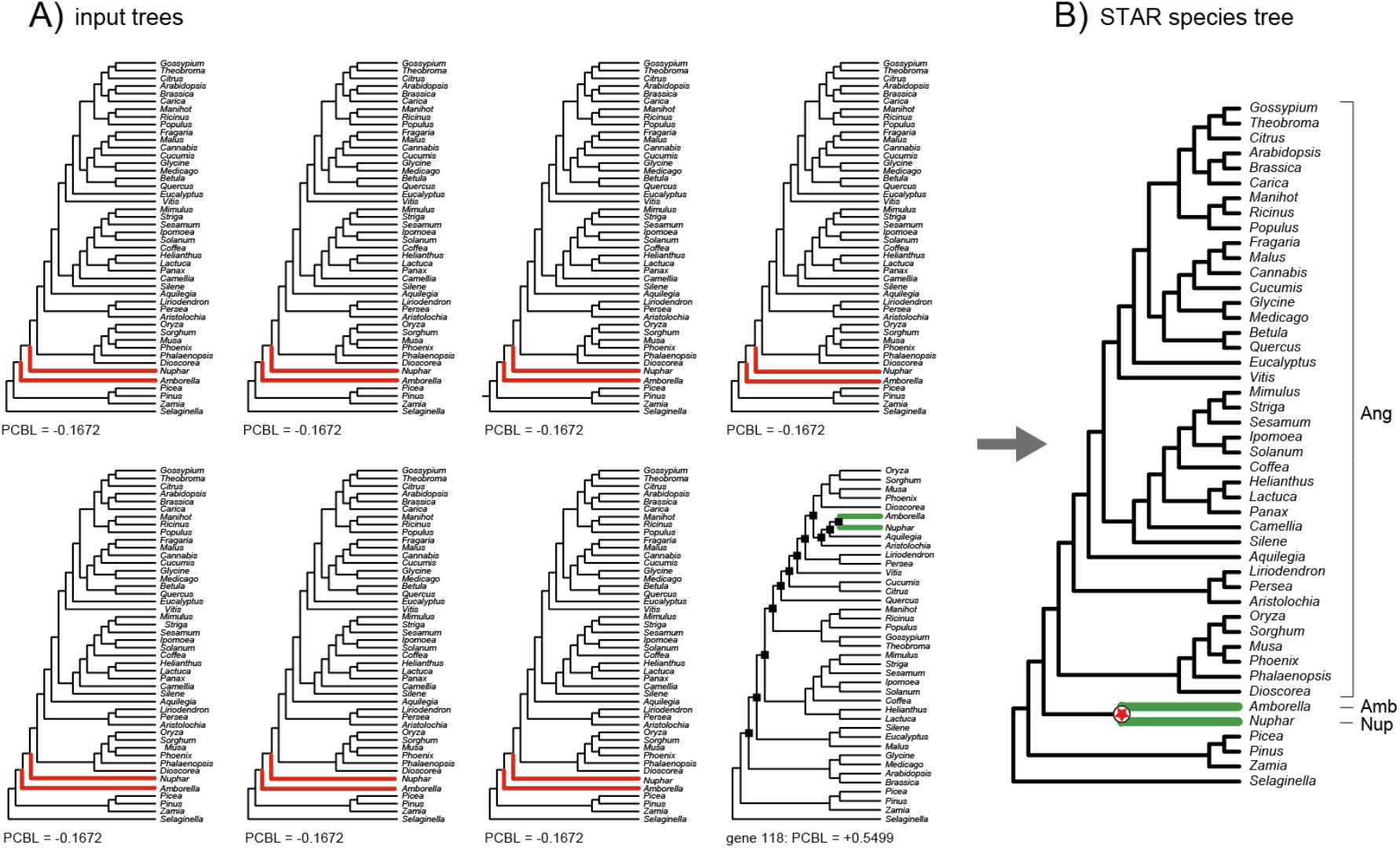
A hypothetical example that demonstrates the extreme impact of a single badly misrooted gene tree (A) for inference of a distance-based species tree (B). STAR analysis of seven identical topologies and a single misrooted gene tree yielded a species tree that supports the conflicting clade resolved by the single outlier gene tree. The seven identical input topologies match the ASTRAL species tree (Simmons and Gatesy, 2015) for the full angiosperm-plants dataset that positions *Amborella* (Amb) sister to a clade composed of *Nuphar* (Nup) and remaining angiosperms (Ang). The single, misrooted outlier is gene tree #118 from Xi et al. (2014) in which *Amborella* groups with *Nuphar* (Nup), and these two taxa are nested 12 nodes within angiosperms (black squares at nodes). The red star (B) marks the single clade in the inferred STAR species tree that conflicts with each of the seven identical input trees (A). STAR PCBL scores for input trees are shown.

Despite the very different underpinnings of distance-based coalescent methods and MP-EST, the latter suffers from similar biases (Gatesy et al., 2019). Many of the same ‘misrooted’, ‘apically nested’ gene trees provided high support for the *Amborella* + *Nuphar* clade in MP-EST analysis. By contrast, ASTRAL results agreed with ML concatenation in contradicting the *Amborella* + *Nuphar* clade (Fig. 2D). We therefore suggest that this congruent high support in MP-EST, STAR, and NJst analyses is due to similar biases of all three methods. Likewise, congruent support in STAR, NJst, and MP-EST analyses of the spider and skink datasets was contradicted by both ASTRAL and ML concatenation results (Fig. 2A, C). Based on PCBL (Figs. 4-5) and PCS (Gatesy et al., 2019), we again contend that an ‘apical nesting’ bias in STAR, NJst, and MP-EST generated congruence that should not be trusted in phylogenomic analyses of these datasets.

#### 3.2.3 ‘Basal dragdown’ bias in distance-based coalescent methods

In outlier gene trees with highest PCBL for the mammal dataset (Liu et al., 2017a), the aye-aye, *Daubentonia* (Chiromyiformes), is widely separated from the dubious *Otolemur* (Lorisiformes) + *Microcebus* (Lemuriformes) clade by 26 to 34 incongruent nodes (Fig. 8). These highly incongruent gene trees presumably ‘dragged’ the divergence of *Daubentonia* to a more basal position in distance-based species tree, away from *Daubentonia*’s closest relative, *Microcebus* (Fabre et al., 2009; Perelman et al., 2011; McLain et al., 2012; Springer et al., 2012; Finstermeier et al., 2013; Marciniak et al. 2021). To quantify the severity of this problematic ‘basal dragdown’ bias, we analyzed several contrived datasets.

Figure 11 shows the first hypothetical case in which the strongest outlier gene tree (#878) with highest STAR and NJst PCBL for the *Otolemur* + *Microcebus* clade (Fig. 7A-B) was combined with multiple, perfectly congruent topologies that match the ASTRAL species tree for the full mammal dataset that instead supported *Daubentonia* + *Microcebus* (Fig. 2E). When the single outlier gene tree was combined with 17 identical trees that include the *Daubentonia* + *Microcebus* clade, the *Otolemur* + *Microcebus* clade that is contradicted by 17 of 18 (94%) input trees was supported by STAR analysis. The *Daubentonia* divergence was ‘dragged’ one node more basal in the species tree relative to its position in the 17 congruent input trees (Fig. 11). For NJst, results were the same. In this hypothetical situation, ASTRAL did not suffer from the same ‘basal dragdown’ bias that characterized both distance-based coalescent methods. Just a 2:1 ratio of input trees yielded an ASTRAL species tree that supports the *Daubentonia* + *Microcebus* clade, as expected in the absence of bias.

**Figure 11.**
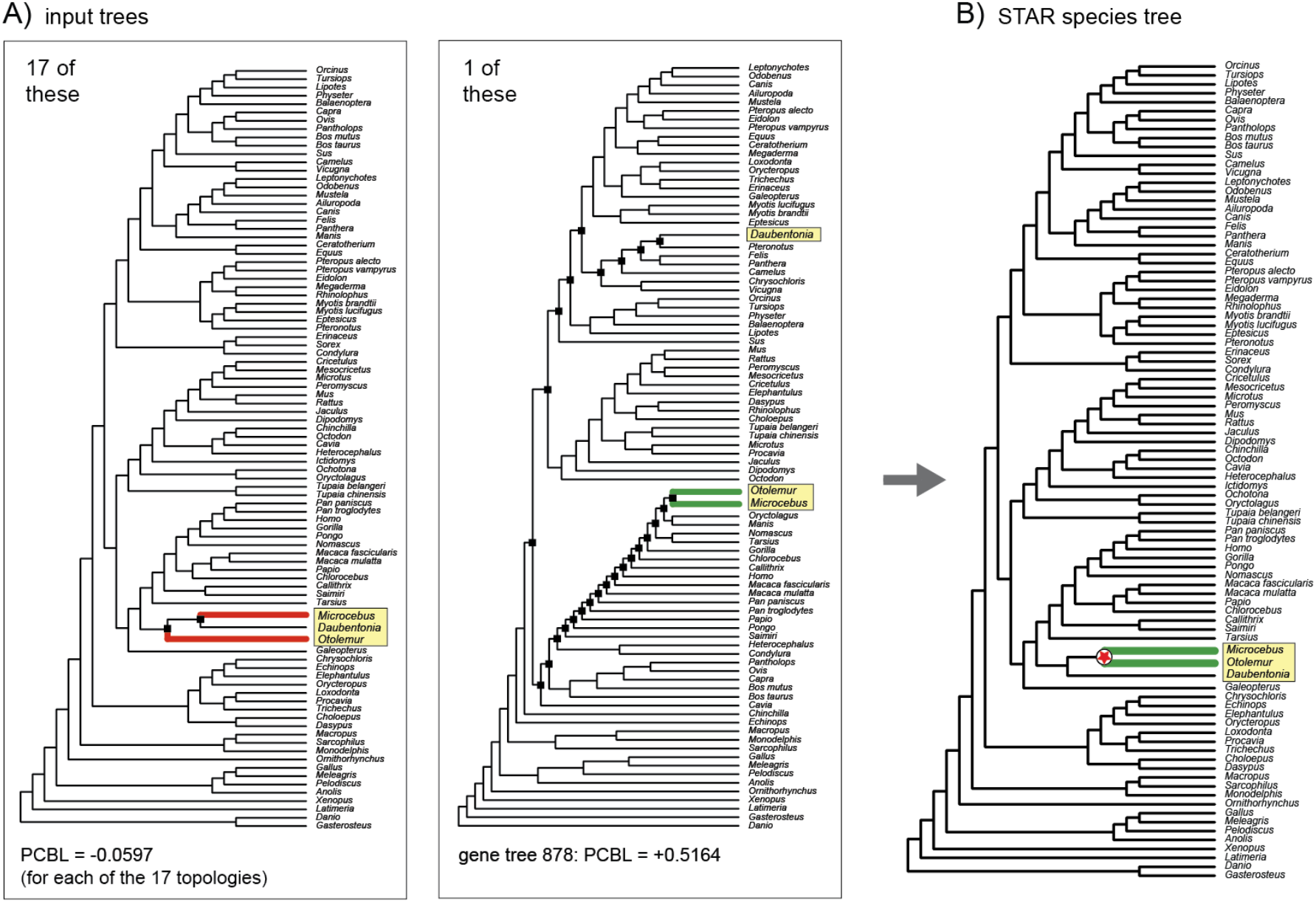
A hypothetical example that demonstrates the extreme impact of a single outlier gene tree (A) for inference of a distance-based species tree (B). STAR analysis of 17 identical topologies and a single outlier gene tree yields a species tree that supports the conflicting clade resolved by the single outlier. The 17 identical input topologies match the ASTRAL species tree for the full mammal dataset of Liu et al. (2017a) that resolves a *Daubentonia* (Chiromyiformes) + *Microcebus* (Lemuriformes) clade. The single outlier is gene tree #878 in which *Microcebus* + *Otolomer* (Lorisiformes) is resolved, and these two strepsirrhine primates are widely separated (black squares at nodes) from the third strepsirrhine (*Daubentonia*). The red star (B) marks the single clade in the inferred STAR species tree that conflicts with each of the 17 identical input trees (A). STAR PCBL scores for input trees are shown.

A second hypothetical case based on the mammal dataset further demonstrates that PCBL is especially useful for identifying outlier gene trees that strongly impact distance-based coalescent results due to the ‘basal dragdown’ bias. Figure 12 illustrates a scenario in which the gene tree with the second highest STAR PCBL score (#86) at the *Otolemur* + *Microcebus* node (Fig. 7A-B) was combined with multiple, perfectly congruent topologies that match the ASTRAL species tree for the full mammal dataset (Fig. 2E). Surprisingly, the outlier gene tree #86 does not resolve *Otolemur* + *Microcebus*, even though PCBL was extremely high for this clade (Fig. 7A-B). When the single outlier was combined with 11 identical gene trees that include the *Daubentonia* + *Microcebus* clade, the *Otolemur* + *Microcebus* clade that is contradicted by 12 of 12 (100%) input trees was supported by STAR analysis (Fig. 12). In this example, support for the dubious *Otolemur* + *Microcebus* clade was completely emergent (i.e., ‘hidden coalescent support’ of Gatesy and Springer [2014]) as was support for two additional emergent clades (*Lipotes* + *Physeter*; *Elephantulus* + *Chrysochloris*). For NJst, results are the same as for STAR, but in this contrived scenario, ASTRAL again did not suffer from a ‘basal dragdown’ bias.

**Figure 12.**
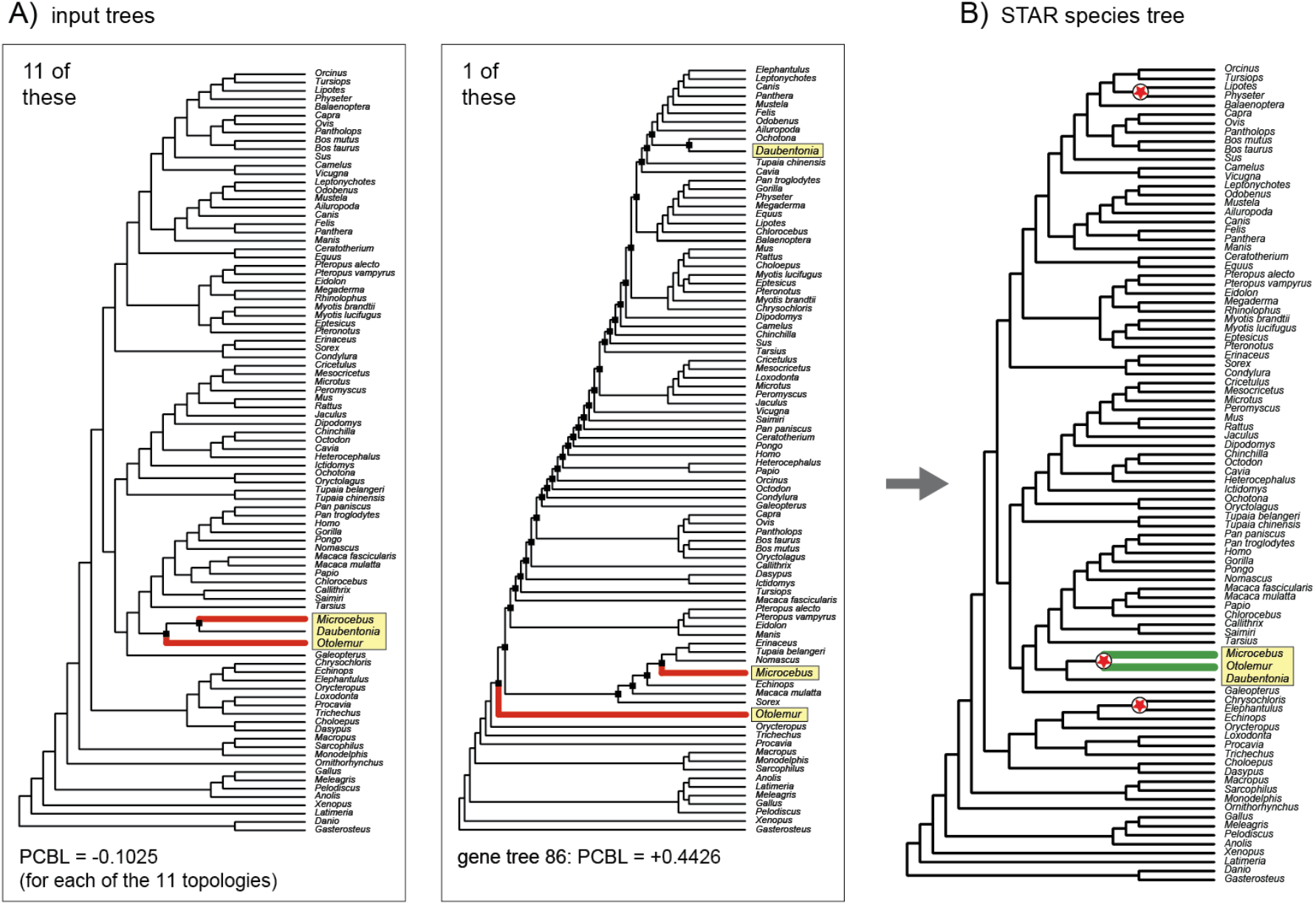
A hypothetical example of extreme hidden support that demonstrates the large impact of a single outlier gene tree (A) for inference of a distance-based species tree (B). STAR analysis of 11 identical topologies and a single outlier gene tree yields a species tree that supports a dubious *Microcebus* + *Otolomer* clade that is contradicted by all 12 input trees. Additionally, two other emergent clades that are resolved in none of the 12 input trees are supported by STAR analysis (red stars). The 11 perfectly congruent trees match the ASTRAL species tree for the full mammal dataset of Liu et al. (2017a) that resolves a *Daubentonia* (Chiromyiformes) + *Microcebus* (Lemuriformes) clade. The single outlier is gene tree #86 in which Strepsirrhini is polyphyletic (genera enclosed in yellow rectangles) and *Daubentonia* is widely separated from the other two strepsirrhines by many internal nodes (black squares at nodes). STAR PCBL scores for input trees are shown.

Like the ‘missing taxa’ bias and the related ‘misrooting’ and ‘apical nesting’ biases, a ‘basal dragdown’ bias already has been documented for MP-EST (Gatesy et al., 2017, 2019). Congruence between distance-based coalescent methods (STAR, NJst) and MP-EST therefore should be treated with caution when conflicts are observed relative to phylogenetic methods that are not known to have the ‘basal dragdown’ bias (e.g., ASTRAL and ML concatenation). This is precisely the case for the *Otolemur* + *Microcebus* clade which was supported by NJst, STAR, and also MP-EST (this study; MP-EST v.1.5 with 1000 independent search replicates) but was contradicted by both ASTRAL and ML concatenation (Fig. 2E).

### 3.3 Implications of biases for interpretation of simulation results

A general pattern that emerged in this study is that outlier gene trees with many taxa (and therefore many internodes) can bias distance-based species trees the most (Figs. 9-12, S2). This is why initial simulations that tested the relative accuracy and efficiency of summary coalescent methods were insufficient (Gatesy and Springer, 2014). Early simulations using MP-EST, STAR, and NJst included just a few taxa (e.g., Liu et al., 2009b, 2010), and it is impossible to generate highly biased, outlier gene trees that conflict at numerous internodes (e.g., Fig. 8) when all simulated gene trees are small.

Limited simulations with few taxa therefore should not be interpreted as general predictors of a phylogenomic method’s utility when the simulations do not cover areas of the overall ‘tree-space’ where damaging biases of particular methods can occur. For example, Bayzid and Warnow (2013) simulated more deeply diverging species trees with 11-17 taxa for 5-100 genes and found that MP-EST was uniformly bested by ML concatenation. Subsequently, Mirarab and Warnow (2015) simulated still larger datasets. ML concatenation generally outperformed MP-EST, NJst, and often ASTRAL for large species trees (100-500 taxa) with relatively deep divergences (2M-10M generations) when 1000 loci were sampled (their figure 2), presumably due to high reconstruction error in large gene trees (their figure 5). These results contrasted with earlier simulation work based on small species trees with shallow divergences, wherein multiple hits at nucleotide sites are rare (e.g., Kubatko et al., 2009; Liu and Edwards, 2009; Liu et al., 2009b, 2010; Liu and Yu, 2011; Hovmöller et al., 2013).

Detailed interrogation of empirical data (e.g., Brown and Thomson, 2017; Shen et al., 2017, 2021), as well as hypothetical scenarios based on empirical datasets (e.g., Figs. 9-12) can provide compelling insights limitations of different phylogenomic methods and complement traditional simulation work. In particular, strong conflicts among methods observed in empirical systematic work are useful for determining which simulation conditions should be explored to more fully understand biases revealed by partitioned support indices. Our results (Gatesy et al., 2017, 2019; Simmons et al., 2022; this study) specifically indicate that large trees with many deeply divergent taxa are challenging for MP-EST, STAR, and NJst, and we predict that the biased support of individual outlier gene trees will become even more problematic as taxon sampling is augmented still further. Given that increased taxon sampling can improve phylogenetic accuracy of gene trees (Hedtke and Hillis, 2006), extensive missing sequence data that are unequally distributed among taxa (Zhong et al. 2013; Hosner et al., 2016; Liu et al., 2017a; Oliveros et al., 2019), long outgroup branches (Xi et al., 2014; Cloutier et al., 2019), unequal rates of evolution among lineages (e.g., Song et al., 2012; Esselstyn et al., 2017), and rapid divergences that are deep in time (Chiari et al., 2012; Linkem et al., 2016) should be simulated in large trees to assess the relative success of phylogenomic algorithms in these contexts.

### 3.4 Should certain ‘shortcut’ coalescent methods be retired?

Contra Liu et al. (2019; p. 277), we do not think that summary coalescent methods are generally “less susceptible to error due to deviations of single genes from neutral expectations”. Indeed, by partitioning the support/influence of individual gene trees in MP-EST, STAR, and NJst analyses (Gatesy et al., 2017, 2019; Simmons et al., 2022; this study), it is evident that outlier gene trees that contradict the clade of interest commonly have the most impact for the resolution of that clade (Figs. 4-12), and this biased influence can explain robustly supported conflicts among different coalescent methods and between coalescent and concatenation results. In some cases, the deletion of just a few outliers can overturn clades that had high bootstrap support. Worse, the biases of STAR, NJst, and MP-EST are shared biases that can lead to congruent misplacements of taxa that uniformly conflict with more robust methods (Fig. 2).

The extreme overweighting of inaccurately reconstructed gene trees is a profound problem (e.g., Figs. 9-11), in particular if a researcher is attempting to resolve challenging systematic relationships in the Tree of Life, such as rapid radiations that are deep in time. There is a hope that the analysis of genome-scale data using ILS-aware methods may finally answer such questions (Edwards et al., 2016; Liu et al., 2019), but at short internodes, support for the correct resolution versus alternatives can be nearly equal (e.g., Cloutier et al., 2019). In this context, individual outlier gene trees that are overweighted 7×, 11×, or 17× (Figs. 10-12) more than they should be are especially damaging, and this problem is compounded by missing taxa in gene trees (Figs. 9, S2; Gatesy et al., 2019).

An important question to ask at this point is whether phylogenomic coalescent methods with shared biases should be retired, given that alternative summary coalescent methods, such as ASTRAL, are less biased. Based on our results, we see no reason to continue using STAR, NJst, and MP-EST as alternatives or supplements to ASTRAL. A more tempered response would be to proceed cautiously when applying biased summary coalescent methods, in particular under conditions that can accentuate these problems (i.e., large gene trees, outgroup misrooting, nonrandomly distributed missing data, extensive gene tree reconstruction errors). Numerous published studies that relied upon STAR, NJst, and/or MP-EST might be tainted by the biases characterized here, and where conflicts with more robust phylogenomic methods have been noted (e.g., McCormack et al., 2012; Song et al., 2012; McLean et al., 2019; Meiklejohn et al., 2016; Cloutier et al., 2019), partitioned support indices would likely help to arbitrate such conflicts.

## 4. Conclusions

The ten main conclusions of our study are as follows:

1. PCBL (Fig. 1) quantifies the influence of each gene tree at every internode supported by a distance-based summary coalescent method.
2. Scaled PCBL (Figs. 1, S2) summarizes the *proportional* influence of each gene tree relative to the total branch length that subtends a clade.
3. Automation of PCBL permits rapid calculation of PCBL scores for genome-scale datasets of hundreds to thousands of gene trees using either STAR or NJst (https://github.com/dbsloan).
4. Application of PCBL to five empirical phylogenomic datasets showed that in four cases (Zhong et al., 2013; Bond et al., 2014; Xi et al., 2014; Linkem et al., 2016), contentious relationships that were central to the conclusions of the original studies were sensitive to the removal of just one to nine gene trees with high PCBL scores (Figs. 3-6). In some cases, clades with high bootstrap support for distance-based methods (Fig 2B-D) flipped to alternative phylogenomic resolutions after removal of over-weighted gene trees that were identified by PCBL (Figs. 4-6).
5. PCBL scores quantified the overweighting of outlier gene trees in distance-based coalescent analyses. Gene trees that did not even resolve the clade of interest often had extremely high PCBL scores for that clade (Figs. 3C, 4C, 5C, 6C, 8), which indicated the strongly biased influence of these outliers (e.g., Fig. 12).
6. PCBL scores quantified the striking impact of missing data in distance-based species trees, including both missing taxa in gene trees (Figs. 3, 9) as well as taxa with partial sequences (Figs. 4, 8, S3, S4). The ‘missing taxa’ bias that was previously characterized in MP-EST and ASTRAL analyses also applied to the distance-based coalescent methods STAR and NJst (Fig. S2).
7. ‘Misrooting’ (Simmons and Gatesy, 2015; Simmons et al., 2022), ‘basal dragdown’ (Gatesy et al., 2017, 2019), and ‘apical-nesting’ biases (Gatesy et al., 2019) that were previously characterized for MP-EST also applied to STAR and NJst (Figs. 4-7).
8. Shared biases across different summary coalescent methods indicate that extreme caution should be exercised when interpreting congruence among these methods relative to conflicts with topologies supported by concatenation and other systematic approaches.
9. Partitioned support indices, including PCBL, can now be used to better understand strong conflicts between results supported by alternative phylogenomic inference methods, including ASTRAL, MP-EST, STAR, NJst, ML concatenation, Bayesian concatenation, and parsimony concatenation.
10. The general PCBL approach can be extended to additional distance-based coalescent methods, such as STEAC or ASTRID, to better understand the impacts of missing data, outlier gene trees, and other factors in phylogenomic analyses using these approaches.

## Abbreviations

ASTRAL: accurate species tree algorithm
ASTRID: accurate species trees from internode distances
bp: base pair
ILS: incomplete lineage sorting
ML: maximum likelihood
MP-EST: maximum pseudo-likelihood for estimating species trees
NJst: neighbor joining species tree
PCS: partitioned coalescence support
PP: posterior probability
STAR: species tree estimation using average ranks of coalescences
STEAC: species tree estimation using average coalescence times
UCE: ultraconserved element

## 5. Acknowledgements

We thank M. Hedin and R. Baker for helpful comments on various sections of the manuscript. This research was funded by NSF grant DEB-1457735 (J.G. and M.S.S.), NSF grants IOS-1829176 and IOS-2114641 (D.B.S), and an NSF-funded GAUSSI Graduate Fellowship (DGE-1450032) to J.M.W.

## 7. Figures

**Figure S1.**
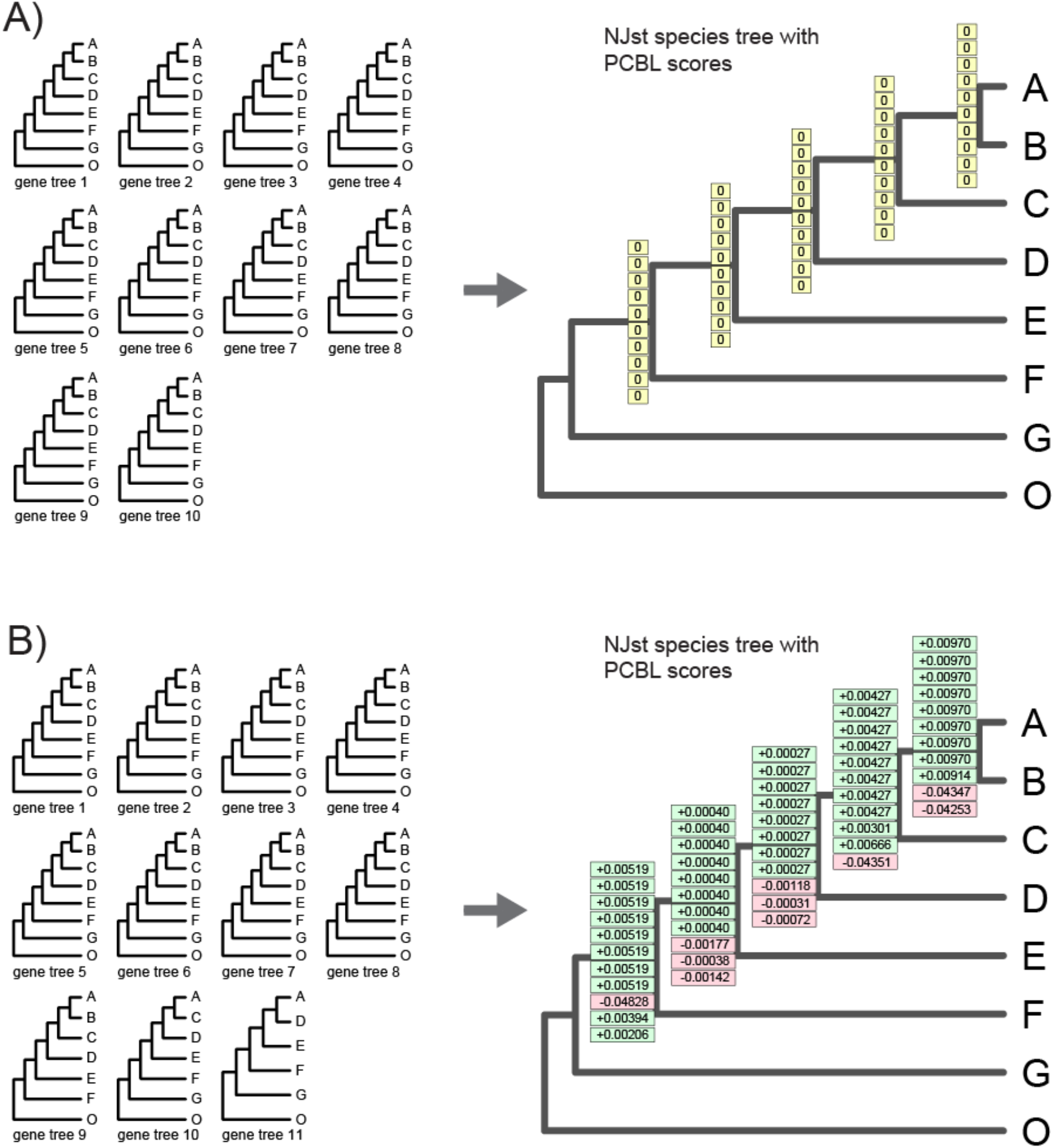
Hypothetical examples that show NJst PCBL scores for cases in which there is complete congruence among gene trees either without (A) or with (B) missing taxa in some input trees. For example (A), all ten gene trees include all eight taxa. NJst PCBL is zero for every gene at every internal node, because removal of any single gene tree from analysis does not impact the average internodal distance between any pair of taxa in gene trees. For example (B), some taxa are missing from three of the 11 gene trees (#9, #10, #11). Gene trees with missing taxa reduce the average internodal distances between some pairs of taxa relative to a complete sampling of taxa for all gene trees. PCBL scores are therefore slightly positive at all nodes for complete gene trees, while gene trees with missing taxa have negative PCBL at some internal nodes.

**Figure S2.**
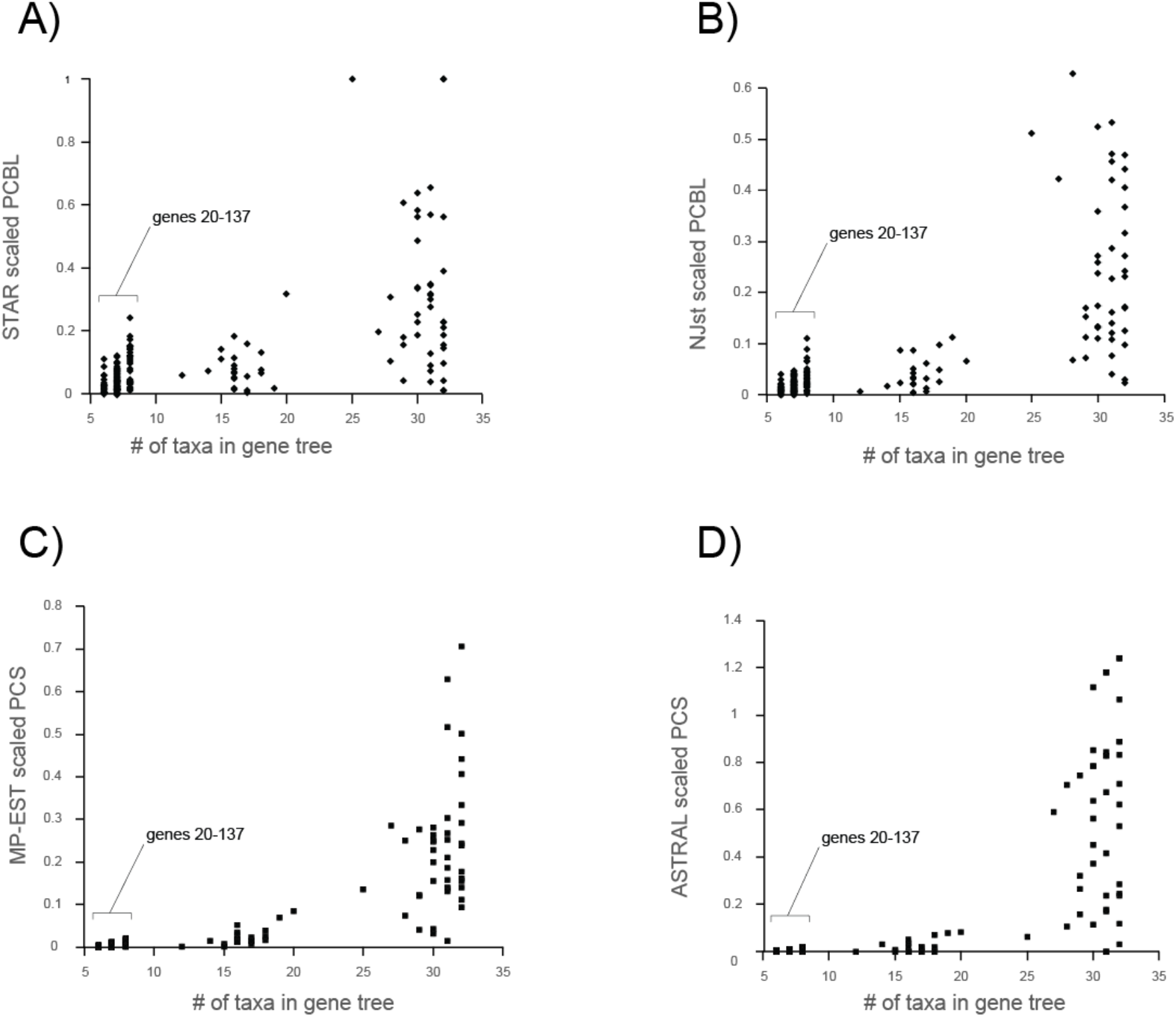
Gene tree size (number of taxa) versus the absolute value of STAR scaled PCBL (A), NJst scaled PCBL (B), MP-EST scaled PCS (C), and ASTRAL scaled PCS (D) for the green-plants dataset of 184 gene trees (Zhong et al., 2013). The absolute value of scaled partitioned support (Y axis) is plotted against gene tree size (X axis). Note that the scaled partitioned supports for small gene trees are consistently low, but scaled partitioned supports for larger gene trees extend across a much broader range of values. The set of 118 small gene trees (each 6-8 taxa; gene trees #20-137) from Zhong et al. (2013) that have little influence in STAR, NJst, MP-EST, and ASTRAL analyses are bracketed.

**Figure S3.**
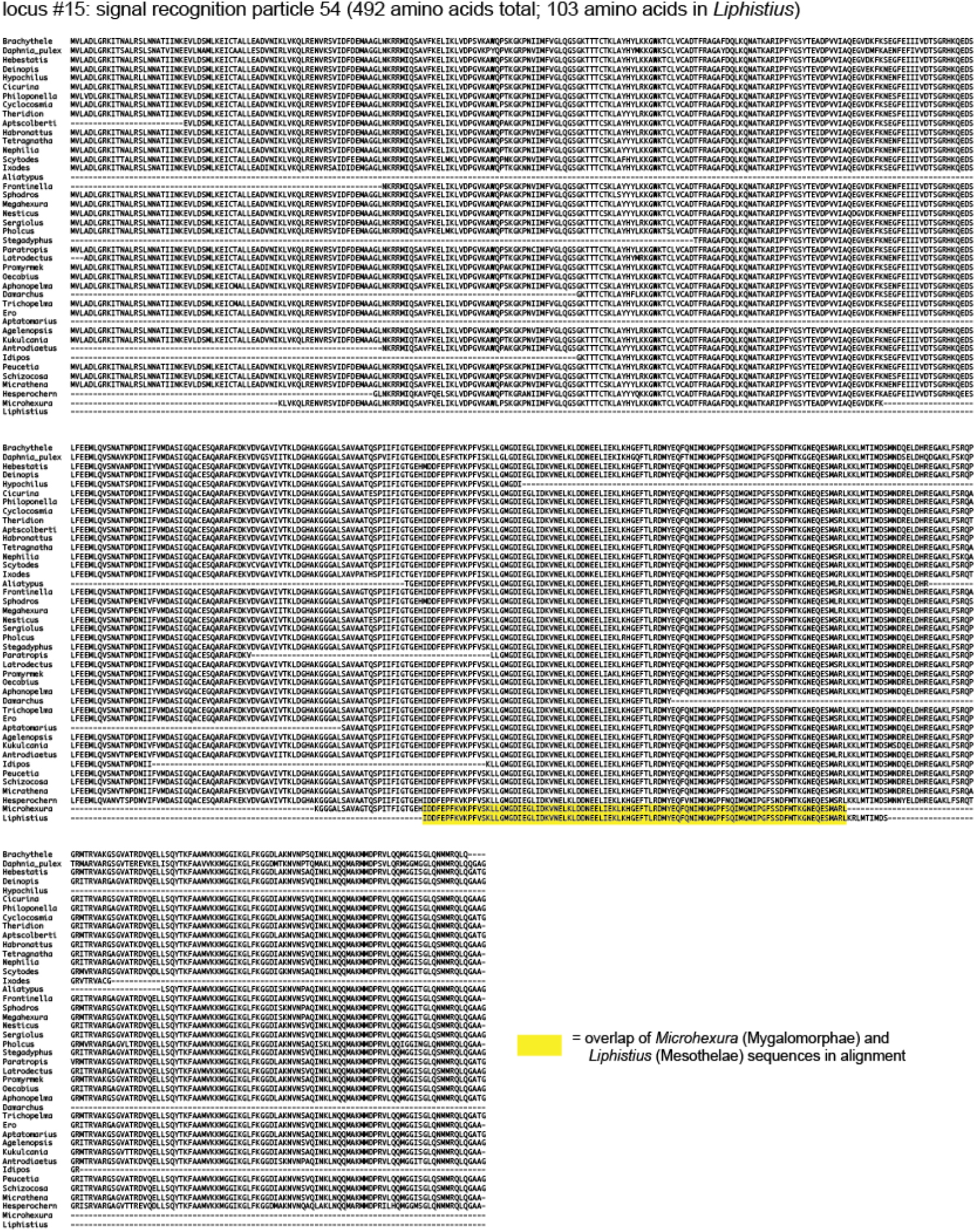
Amino acid sequence alignment for locus #15 (signal recognition particle 54) from Bond et al. (2014). Note the extensive missing data for *Liphistius* (Mesothelae) and *Microhexura* (Mygalomorphae). The amino acid sequences for these two distantly related taxa (yellow) are identical where they overlap.

**Figure S4.**
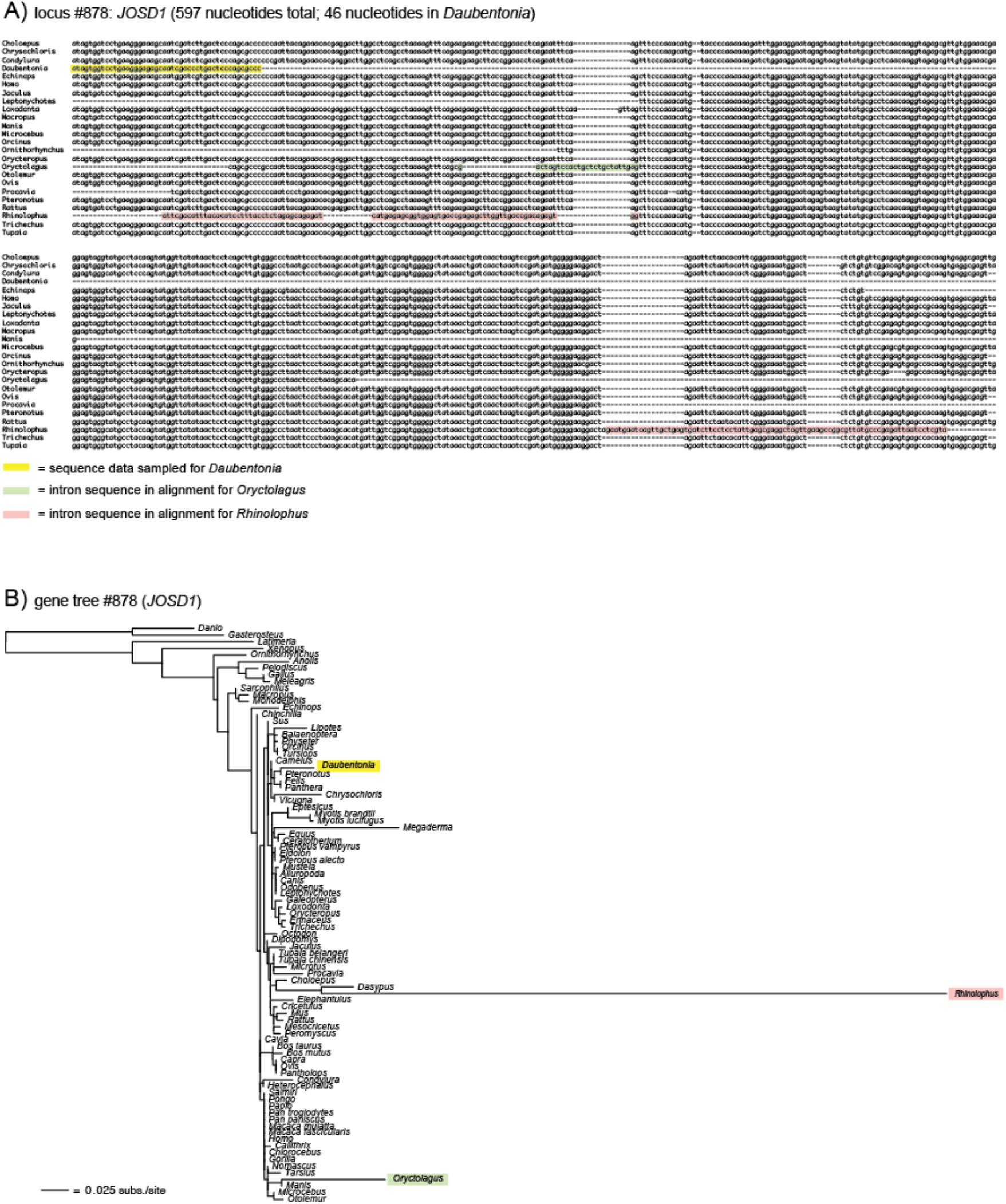
Sequence alignment (A) and ML phylogram (B) for locus #878 (*JOSD1*) from Liu et al. (2017a). A subset of taxa from the full alignment of 87 species is shown. Note the extensive missing data for the aye-aye (*Daubentonia*, Chiromyiformes). The partial sequence for *Daubentonia* (46 bp; yellow highlighting) drives misplacement of this species in the *JOSD1* gene tree. Given extensive missing data, the *Daubentonia* sequence cannot be distinguished from three distantly related mammalian taxa in the alignment: a rabbit (*Oryctolagus*), a seal (*Leptonychotes*), and platypus (*Ornithorhynchus*). The extremely long terminal branch in the phylogram (B) is due to misannotation and subsequent misalignment of *JOSD1* sequences from the bat *Rhinolophus* (pink) in the multispecies alignment (A). In the first block of sequences, *JOSD1* intron 1 from *Rhinolophus* is aligned to exon 1 in other species, and in the second block, *Rhinolophus* intron 3 is aligned to exon 4. There are additional homology errors in sequences for *Oryctolagus* (green) and *Megaderma* (not shown in alignment) that are due to further misannotations. For *Oryctolagus, JOSD1* intron 1 is misaligned to exon 1 in the other species. For *Megaderma*, the 3’ untranslated region of *JOSD1* and sequences from the *TOMM22* gene are inappropriately aligned to *JOSD1* exon 4 in the other species. Such homology errors in the protein-coding-sequence alignments of Liu et al. (2017a) are widespread in their overall mammal dataset (see Gatesy and Springer, 2017).

